# A revision of the African *Russula radicans* and allies in subgen. *Heterophyllidiae* provides an example of a clade that exhibits recent diversification and extensive phenotypic plasticity

**DOI:** 10.1101/2024.11.10.622834

**Authors:** B Buyck, TW Henkel, C Manz, SQ Cao, M Amalfi, XH Wang

**Affiliations:** Institut de Systématique, Evolution, Biodiversité (ISYEB), Muséum national d’Histoire naturelle, CNRS, Sorbonne Université, EPHE, Université des Antilles, CP 39, 57 rue Cuvier, 75005 Paris, France; Department of Biological Sciences, Humboldt State University, Arcata, California 95521, Unites States of America; Department of Biological Sciences, Goethe University, Frankfurt am Main, Germany; Meise Botanic Garden, Nieuwelaan 38, 1860 Meise, Belgium - Fédération Wallonie– Bruxelles, Service Général de l’Enseignement Universitaire et de la Recherche Scientifique, Brussels, Belgium; CAS Key Laboratory for Plant Diversity and Biogeography of East Asia, Kunming Institute of Botany, Chinese Academy of Sciences, Kunming 650201, P. R. China

**Keywords:** African tropics, ectomycorrhizal, Fabaceae, infrageneric classification, Madagascar, Russulaceae, *Uapaca*

## Abstract

*Russula* subsection *Radicantinae* is described for *R. radicans* and allies, while subsection *Aureotactinae* is again restricted to *R. aureotacta*, the subsectional type species that has never been recollected since its original description. *Russula radicans,* originally described from Madagascar, as well as the Central African *R. brunneoannulata,* are epitypified. *Russula xylophila* is the first confirmed annulate species belonging to Sect. *Ingratae* and a close relative to *R. oleifera.* An ITS sequence has been obtained from the *R. xylophila* holotype collection made in 1928, but it is slightly different from those obtained from all new collections. For this reason and some minor morphological differences, we refrained from epitypification. *Russula acriannulata*, another species attributed to *Aureotactinae* in the past, is now also excluded from the subsection, but needs more sequencing to solve its systematic placement. The Malagasy *Russula cibaensis* and *R. tapiae*, as well as the Central African *R. cameroonensis* and *R. afrovinacea* are new species in *Radicantinae*.At least three more undescribed species are revealed by environmental sequences. The West African *R. sankarae* is reduced to a form of *R. radicans*. The latter species exhibits an impressive phenotypic plasticity in response to differences in habitat or host associations. It has a very wide distribution covering littoral rain forest in eastern Madagascar (forma *radicans*), Zambezian woodlands in Central and Eastern Africa (forma *miomboensis* fo. nov.) and gallery forests in West Africa (forma *sankarae*).

## INTRODUCTION

In his monograph on the Russulaceae of Madagascar, Heim (1938) introduced several new infrageneric taxa in the genus *Russula* to accommodate new, morphologically unique species. One of the more interesting examples of these unique morphologies was *R. radicans* Heim, a strongly yellowing, slender, extremely thin-fleshed and annulate species with a radicating stipe. Heim placed it in his “*Pelliculariae”* Heim *nom. inval*. (unranked, but likely equivalent to a new subgenus), together with additional species producing unusually thin-fleshed and fragile basidiomata. Within *Pelliculariae*, Heim recognized again two unranked groups (to which he refers as sections): (1) *Aureotactae* Heim *nom. inval*. for strongly yellowing species, and (2) *Heliochromae* Heim *nom. inval*. for those lacking the yellowing context. *Russula radicans* was thus a member of *Aureotactae,* the type species of which was *R. aureotacta* Heim, the single other strongly yellowing species he described from Madagascar. The latter species differed principally from *R. radicans* in the absence of both an annulus and a radicating stipe. Heim interpreted the sometimes lignicolous fruiting habit and radicating stipe of *R. radicans* as indications for a possibly saprotrophic lifestyle, an exception in a genus that was assumed to be strictly ectomycorrhizal. This hypothetical saprotrophy convinced Heim to place *R. radicans* in a subsection of its own, *Radicantes* Heim *nom. inval*., thereby leaving the single other strongly yellowing species, *R. aureotacta,* as sole member of an equally monotypic subsection *Aureotactinae* Heim. *nom. inval*. Some thirty years later, Heim (1970) placed a second, small and thin-fleshed but pink coloured species, *R. mimetica* Heim, in his *Radicantes*. This placement was essentially based on the more or less radicating stipe, presence of a ring and similar spore ornamentation, notwithstanding the absence of a yellowing context. On the basis of the different microscopic features (particularly the absence of cystidioid hyphae throughout the tissues) and the absence of a distinct yellow discoloration of the fruiting bodies, Buyck (1989b) transferred *R. mimetica* to subgenus *Heterophyllidiae,* where he placed it in a subsection of its own, *Mimeticinae* Buyck, together with some additional South American and tropical African species.

Since the early studies by Heim, neither *R. radicans* nor *R. aureotacta* have been discussed again. In his early studies, Heim (1937, 1938a, 1943, 1970) extensively addressed the unexpected presence of a distinct annulus in some of these tropical *Russula* species. Later, however, the annulate condition turned out to be a rather frequent phenomenon among tropical African members of *Russula* subg. *Heterophyllidiae* sensu Buyck et al. (1994, 2018). Heim demonstrated that this annulus originated as an extension of the pileipellis protruding beyond the young pileus margin until it finally made contact with the stipe; the subsequent expansion of the pileus would then rupture the pileipellis at the point where the underlying pileus trama ceases, leaving thus a fringe of pileipellis tissue hanging around the stipe. Being not convinced of the taxonomic value of a very thin context or the presence of an annulus in the delimitation of subgeneric ranks, Buyck (1989, 1992) abandoned the concept of *Pelliculariae* and transferred most of its species to subg. *Heterophyllidiae* based on shared similarities in micromorphology of the pileipellis and basidiospores.

With the names of both of Heim’s subsections, *Aureotactinae* and *Radicantes,* being invalid, Buyck (1990) validated subsect. *Aureotactinae* rather than *Radicantes*, thereby putting the accent on the shared yellowing context as an important character, rather than on the assumption that the lignicolous habit and associated radicating stipe were an indication of saprotrophy (see also Buyck & Horak, 1999). *Russula radicans* was thereby transferred to *Aureotactinae*. Among the many similarities shared between *R. radicans* and *R. aureotacta,* Heim cited a similar overall basidioma size, yellow-orange colours, strongly yellowing and thin context, whitish spore print, peeling cuticle, strongly striate-tuberculate pileus margin, lacunar stipe, insensitivity to FeSO_4_ and similar type of basidiospore ornamentation. To our surprise, however, Heim (1938) considered pileocystidia to be absent in both species (illustrated in Heim 1938, fig. 46), notwithstanding the fact that these are very abundant, conspicuous in size and filled with abundant and typical, strongly refringent contents.

In his monograph on Central African *Russula* species, Buyck (1989a, 1994) hesitantly placed three additional species in subsect. *Aureotactinae*: two annulate species (*R. brunneoannulata* Buyck and *R. xylophila* Beeli) as well as the exannulate *R. oleifera* Buyck. For none of these three species there was a mention of a yellowing context in the associated field notes, but there seemed at the time no better option to place them considering their microscopic features. Very soon, however, *R. oleifera* was transferred to sect. *Ingratae* based on newly made field observations (Buyck 1992, Härkönen et al. 1993). Around that same period, the annulate *R. acriannulata* Buyck & Härkönen was described as new likely member of *Aureotactinae* (in Härkönen et al. 1993). The latter was a small species collected in dense lower mountain forest at 1800-1900 m alt. in the West Usambara Mountains in Tanzania. Finally, the West African annulate and strongly yellowing *R. sankarae* Sanon & Buyck (in Sanon et al. 2014) was described as a typical member of the subsection.

Whereas all presently known *Aureotactinae* were based on a single or very few specimens collected in the humid forest biome, recent fieldwork has shown that *Aureotactinae* are also common and even locally abundant in the seasonal surrounding woodlands. In this paper, we revisit African and Malagasy *Aureotactinae* on the basis of over 40 newly sequenced African collections representing all of the species that have been assigned to it in the past, with one notable exception: *R. aureotacta* itself, the type species of the subsection, which has yet to be recollected. As discussed below, the absence of new collections for *R. aureotacta* poses problems for the correct interpretation of the subsection of which it is the type species. Therefore, *Aureotactinae* is here restricted again to Heim’s monotypic concept, and a new subsection *Radicantinae* is introduced to harbour *R. radicans* and close relatives. The earliest species are here epitypified and several new species and infraspecific taxa described.

## Materials and methods

### 1. Morphology

All studied collections were photographed in the field or at least in fresh condition and all had accompanying notes and descriptions. Colour references refer to Kornerup & Wanscher (1978). Microscopic characters were examined under a Nikon E400 microscope (Nikon, Tokyo) at a magnification of 1000x. Fragments of hymenium and pileipellis were shortly heated in a 5% KOH solution before observation in Congo red ammonium solution. Basidiospore ornamentation was observed in Melzer’s reagent and gloeocystidia in sulfovanillin (SV). Line drawings were made at a projected magnification of 2400x with the aid of a drawing tube (Y-IDT, Nikon, Tokyo, Japan). Basidiospore dimensions are given following the form (a) b-m-c (d), with m the mean value, b-c containing at least 90% of all values and with extreme values (a, d) enclosed in parentheses. Q indicates the basidiospore length/width ratio.

### 2. Extraction and amplification for Illumina sequencing of the R. xylophila holotype

DNA was extracted using a CTAB method (Doyle and Doyle, 1990) slightly modified in order to obtain DNA with suitable quality from small amounts of material. All specimens were subjected to a set of preliminary standard PCR by using primer pairs ITS1F/ITS4, ITS1F/ITS2 and ITS3/ITS4 (Gardes and Bruns, 1993, White et al., 1990). Successful PCR reactions resulting in a single band observed on a 0.8% agarose gel were purified by adding 1 U of Exonuclease I (EXO) and 0.5 U FastAP Alkaline Phosphatase (SAP, Thermo Scientific, St. Leon-Rot, Germany) and incubating at 37 °C for 1 h, followed by inactivation at 80 °C for 15 min and sent to Macrogen Inc. (Korea and The Netherlands) for sequencing using the same primers as used for PCR. Amplification of the nuclear ribosomal ITS region for the Illumina sequencing was carried out using a two-step PCR process. In the first PCR the regions of interest was amplified using the primers ITS1F/ITS2 and ITS3/ITS4 in order to amplify respectively the ITS1 and ITS2 regions of the rDNA (Gardes and Bruns, 1993, White et al., 1990), while in the second PCR the products of the first amplification were amplified using either ITS5F/ITS2 and ITS3mod/ITS4 tagged primers in order to add different 5 bp identifier tags to distinguish sequences from each specimen (Voyron et al., 2016). In this work 15 different tags were used, and the second PCR was done in four replicates for each couple of tagged primers. The first PCR reaction was carried out in a total volume of 25 μL including 5 μL of 5X Wonder Taq reaction buffer (5 mM dNTPs, 15 mM MgCl2; Euroclone), 0.5 μL of bovine serum albumin (BSA, 10 mg/mL), 0.5 μL each of two primers (10 μM), 0.5 μL of Wonder Taq (5 U/μL), 2 μL of genomic DNA and ddH2O to reach the final volume. The PCR conditions used were: 95◦C for 3 min; 35 cycles of 95◦C for 30 s, 52◦C for 40 s and 72◦C for 45 s; 72◦C for 5 min. The second PCR reaction was performed similarly to the previous amplification except for the absence of the BSA, the use of 2 μL of the first PCR amplicons as template and the use of the tagged primers. The obtained PCR products were then further analyzed on a 5200 Fragment Analyzer System with the dsDNA 915 Reagent Kit (35–5000 bp) (Agilent Technologies, Santa Clara, California, USA). The four replicates of each sample were pooled together and purified with the same EXO-SAP protocol described above. The libraries were sent to IGATech (Udine, Italy) for a paired-end sequencing using the Illumina MiSeq technology (2 × 300 bp). the purified amplicons were mixed in equimolar amount to prepare two sequencing libraries of 15 samples each. The libraries were sent to IGATech (Udine, Italy) for a paired-end sequencing using the Illumina MiSeq technology (2 × 300 bp).

#### NGS Data Processing, OTU Identification and Taxonomic Analysis

Raw data of the Illumina sequencing were analyzed using the Illumina pipeline developed for a community analysis of orchid mycorrhizal soil fungi (Voyron et al., 2016) as guideline. Forward and reverse reads from each library were merged using PEAR v0.9.10 (Zhang et al., 2014) with the quality score threshold set at 28, the minimum length of reads after trimming set at 150 bp and the minimum overlap size set at 100 bp. The merged reads were processed using the software package QIIME v1.9.1 (Caporaso et al., 2010). To avoid incorrect assignment, demultiplexing of sequences was performed with a maximum number of errors in the tag sequence of 0. At the same time, the sequences were processed using a minimum sequence length cutoff of 150 bp, minimum quality score of 28, a sliding window test of quality score of 50, a maximum length of homopolymers of 13, a maximum number of ambiguous bases of 0 and a maximum number of mismatches in forward and reverse primers of 3, and then reoriented when necessary to 5′ to 3′. Sequences were dereplicated with VSEARCH v2.3.4 (Rognes et al., 2016) to form clusters with an identity of 100% and the ITS1 and ITS2 variable regions extracted using ITSx software (http://microbiology.se/software/itsx/; Bengtsson-Palme et al., 2013). The ITSx was used to eliminate the conserved regions where the primers are located as these regions can introduce bias during the subsequent clustering and taxonomic assignment, because they increase similarity among sequences (Bálint et al., 2014). In addition, setting the option -t F (list of organism group Fungi) all the ITS regions not detected as Fungi were eliminated. The picking of the Operational Taxonomic Units (OTUs) was performed with a 98% of similarity clustering using VSEARCH and the clusters with less than 10 sequences were discarded. Chimera sequences were then removed using a de novo chimera detection with UCHIME algorithm (Edgar et al., 2011) implemented in the VSEARCH pipeline. The last release of the UNITE dataset version 7.2 (https://unite.ut.ee) for QIIME was used as reference for the taxonomic assignment of OTUs (Abarenkov et al., 2010; Kõljalg et al., 2013) and, consequently, for the identification of specimen target sequence. The OTU abundance table was created with VSEARCH, considering a 98% of identity, to determine the number of times an OTU was found in each sample pooled in the libraries. The sum of these numbers for a sample gives the total sequences found for that specific sample. Because a number of OTUs returned no matches compared with the UNITE database, we also compared these sequences with those deposited in NCBI GenBank, excluding uncultured/environmental sample sequences, using the BLASTN algorithm (BLAST-search, http://www.ncbi.nlm. nih.gov/BLAST/; Altschul et al., 1997) and the first 10 hits were evaluated considering consistency between hits, E-value, similarity higher than 96%, query coverage, organism source and publication status of the hits.

### 3. Phylogeny

#### Sampling

For a representative sampling of all known subgenera of *Russula,* we basically followed the multi-gene dataset of Buyck et al. (2024) and removed several duplicate samples of the same species. Buyck et al. (2018) and Wang et al. (2019) have shown that *R.* aff. *brunneoannulata* is nested in subgen. *Heterophyllidia.*We sampled all main lineages of the subgenus including the Asian species *R. verrucospora* with isolated position (Song et al. 2018) and the North American *R. redolens.* Sequences were borrowed from Buyck et al. (2018, 2020, 2024), Wang et al. (2019), Rossi et al. (2020) and Cao et al. (2023) or newly generated in this study. To guarantee more solid evidences for phylogenetic inference, we specifically selected samples that could minimize missing data in our multigene phylogeny. In addition, we amplified ITS of 23 specimens of *R. radicans* from four countries, including the epitype from Madagascar, six specimens of *R. sankarae,* including the holotype from Burkina Faso, two specimens of *R. tapiae* sp. nov. and the holotype of *R. cibaensis sp. nov.,* both from Madagascar,, three specimens of *R. xylophila* and three of *R. cameroonensis* sp.nov., both from Cameroon and four specimens, including the epitype, of *R. brunneoannulata* from the Central African Republic. Except for Malagasy *R. aureotacta* and Tanzanian *R. acriannulata,* this sampling covered all known representatives of sect. *Aureotactae*. Newly generated sequences for the five-locus multigene analysis in this study are listed in Table I and in Fig. 1 for the ITS sequences.

**Table 1:**
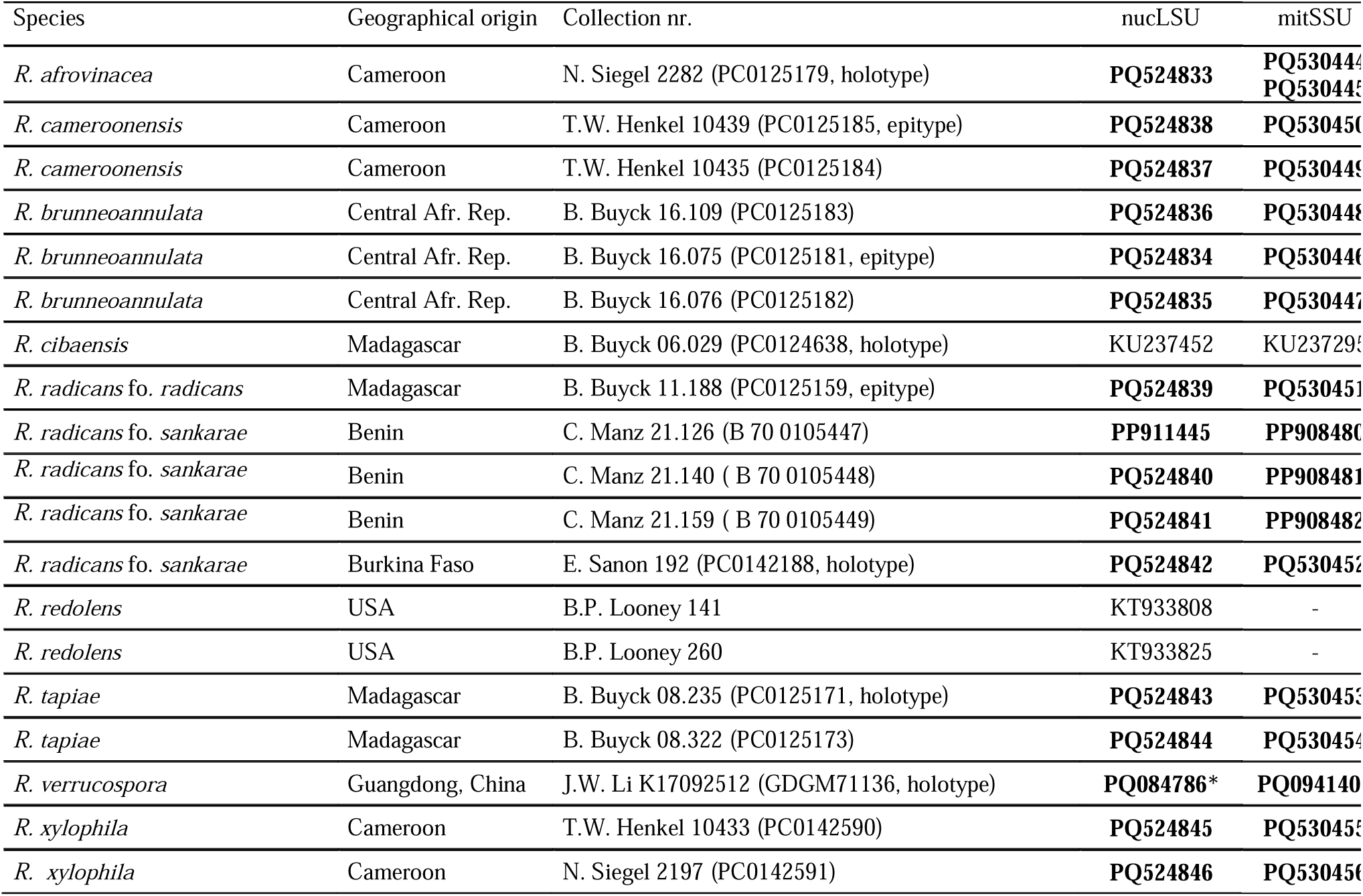
Voucher table for the multigene tree (Fig. 1). Newly produced sequences are in bold.

| Species | Geographical origin | Collection nr. | nucLSU | mitSSU |
| --- | --- | --- | --- | --- |
| *R. afrovinacea* | Cameroon | N. Siegel 2282 (PC0125179, holotype) | **PQ524833** | **PQ530444**<br>**PQ530445** |
| *R. cameroonensis* | Cameroon | T.W. Henkel 10439 (PC0125185, epitype) | **PQ524838** | **PQ530450** |
| *R. cameroonensis* | Cameroon | T.W. Henkel 10435 (PC0125184) | **PQ524837** | **PQ530449** |
| *R. brunneoannulata* | Central Afr. Rep. | B. Buyck 16.109 (PC0125183) | **PQ524836** | **PQ530448** |
| *R. brunneoannulata* | Central Afr. Rep. | B. Buyck 16.075 (PC0125181, epitype) | **PQ524834** | **PQ530446** |
| *R. brunneoannulata* | Central Afr. Rep. | B. Buyck 16.076 (PC0125182) | **PQ524835** | **PQ530447** |
| *R. cibaensis* | Madagascar | B. Buyck 06.029 (PC0124638, holotype) | KU237452 | KU237295 |
| *R. radicans* fo. *radicans* | Madagascar | B. Buyck 11.188 (PC0125159, epitype) | **PQ524839** | **PQ530451** |
| *R. radicans* fo. *sankarae* | Benin | C. Manz 21.126 (B 70 0105447) | **PP911445** | **PP908480** |
| *R. radicans* fo. *sankarae* | Benin | C. Manz 21.140 (B 70 0105448) | **PQ524840** | **PP908481** |
| *R. radicans* fo. *sankarae* | Benin | C. Manz 21.159 (B 70 0105449) | **PQ524841** | **PP908482** |
| *R. radicans* fo. *sankarae* | Burkina Faso | E. Sanon 192 (PC0142188, holotype) | **PQ524842** | **PQ530452** |
| *R. redolens* | USA | B.P. Looney 141 | KT933808 | - |
| *R. redolens* | USA | B.P. Looney 260 | KT933825 | - |
| *R. tapiae* | Madagascar | B. Buyck 08.235 (PC0125171, holotype) | **PQ524843** | **PQ530453** |
| *R. tapiae* | Madagascar | B. Buyck 08.322 (PC0125173) | **PQ524844** | **PQ530454** |
| *R. verrucospora* | Guangdong, China | J.W. Li K17092512 (GDGM71136, holotype) | **PQ084786\*** | **PQ094140** |
| *R. xylophila* | Cameroon | T.W. Henkel 10433 (PC0142590) | **PQ524845** | **PQ530455** |
| *R. xylophila* | Cameroon | N. Siegel 2197 (PC0142591) | **PQ524846** | **PQ530456** |

**Fig. 1.**
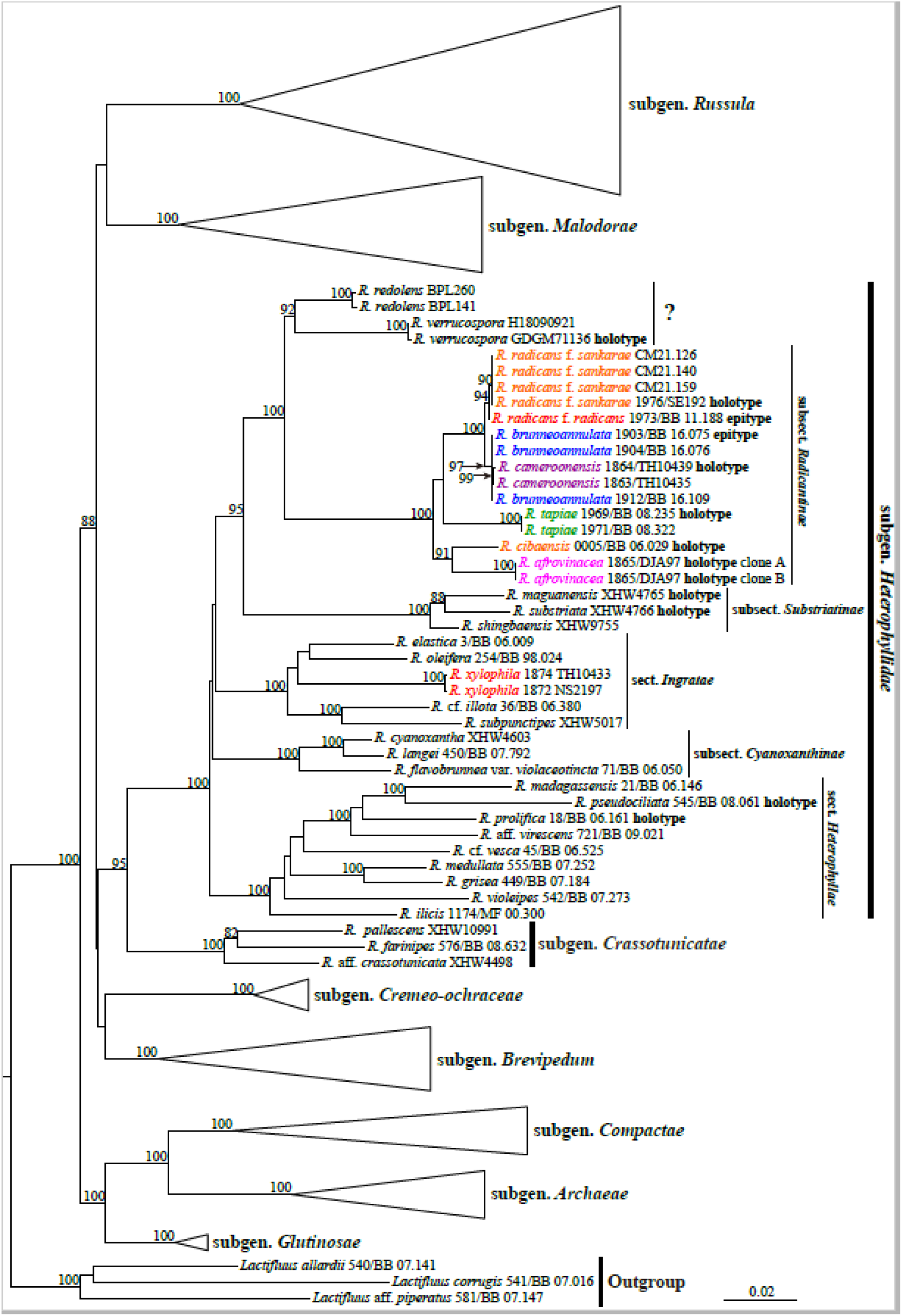
The most likely tree obtained by Maximum Likelihood analysis of the nucLSU-*tef1-rpb2-rpb1*-mtSSU combined dataset, rooted with three species of *Lactifluus,* the sister genus to *Russula.* ML Bootstrap values ≥ 70% are indicating significant support on or by the branches. Infrageneric classification followed Buyck *et al*. (2018, 2024).

#### DNA extraction, PCR amplifications, and sequencing

For DNA extraction, PCR and sequencing, we basically followed the methods and primers of Buyck et al. (2018). For problematic samples, internal primers designed by Cao et al. (2023) were used to separately amplify shorter sections within each locus. We amplified five loci to infer the phylogenetic relationships among the species, following Buyck et al. (2018, 2020): mitochondrial rDNA small subunit (mitSSU), nuclear rDNA large subunit (nucLSU), RNA polymerase II largest (*rpb1*) and second largest subunit (*rpb2*), and translation elongation factor 1-alpha (*tef1*).

PCR amplification and Sanger sequencing for the holotype of *R. verrucospora* (GDGM 71136) failed or partly failed. For this specimen, lower coverage whole genome sequencing or (“genome skimming”) using the next-generation sequencing by Illumina NovoSeq-5500 platform was used to obtain the missing genes. Library preparation followed the protocol described by Zeng et al. (2018). The raw data were de novo assembled using GetOrganelle toolkit (Jin et al., 2020). The target regions were extracted using the reference data from GenBank (MG786052, MK881919, MT085642, MT085569, MT085558 and MK882047).

#### Phylogenetic analyses

Sequences generated by Sanger sequencing were assembled and edited using Sequencher 4.1.4 (Gene Codes Corporation, Ann Arbor, USA). Alignments were performed using ClustalW in BioEdit (Hall 1999) and manually adjusted. Two datasets were used: an ITS dataset to aid species delimitation of *R. radicans* and its allies and a five-locus combined dataset (nucLSU, mitSSU, *rpb1*, *rpb2* and *tef1*) for phylogenetic inference at subgenus level. The ITS alignment was not partitioned. For the five-locus dataset, we followed Buyck et al. (2018) to exclude ambiguous sites and introns and to set the best partitioning, i.e. nucLSU, mitSSU, *rpb1* intron, *rpb1* 1st, *rpb1* 2nd, *rpb1* 3rd, *rpb2* 1st, *rpb1* 2nd, *rpb1* 3rd, *tef1* 1st, *tef1* 2nd and *tef1* 3rd codon positions. The phylogenetic tree for the ITS dataset was rooted using two samples of *R. redolens* and two of *R. verrucospora*, following the relationships inferred by the five-gene dataset. In the analysis of our five-locus dataset, three species of *Lactifluus* were set as outgroups. All best tree searches and bootstrap analyses were conducted in RAxML version 7.7.7 (Stamatakis et al. 2008) using the rapid bootstrap algorithm (RBS). The general time-reversible substitution model with site rate heterogeneity was selected (option -m, GTRGAMMA) and 1000 runs with distinct heuristic starting trees (option -n 1000) were executed. Combinability among the five datasets was examined conducting 1000 RBS heuristics for each single locus dataset. Conflict was assumed when single-locus genealogies for the same set of taxa inferred different relationships (monophyletic versus non-monophyletic) both with significant RBS support values (RBS≥70%; Mason-Gamer and Kellog, 1996). Tree branches were considered significantly supported when RBS values were ≥70% (Alfaro et al. 2003).

## RESULTS

### Phylogeny

Our multigene phylogeny (Fig. 1) retrieved the same topology for subgenera with similar (full) supports as in Buyck et al. (2024). The target samples of *R. radicans* and its allies fell into subgen. *Heterophyllidiae* and formed a long branch receiving full support (MLbs=100%). These samples formed again a fully supported clade with *R. verrucospora* and *R. redolens*. Together with three representatives of subsect. *Substriatinae*, all these species then formed one of the four major clades (MLbs=91%) in the subgenus. *Russula xylophila,* previously considered a close relative to *R. radicans*, grouped with *R. oleifera, R. elastica, R.* cf. *illota* and *R. subpunctipes* in sect. *Ingratae,* another major clade (MLbs=100%) in subgen. *Heterophyllidiae.* The two remaining major clades, both with full support, corresponded to subsect. *Cyanoxanthinae* and sect. *Heterophyllae*. Within subsect. *Radicantinae,* our multigene phylogeny retrieved support for *R. radicans* (MLBs=94%) and its forma *sankarae* (MLBs=88%), for *R. tapiae* (MLBs=100%, which justifies the recognition of *R. cibaensis* as an independent species only known from the holotype, and for *R. afrovinacea* (MLBs=100%) *Russula cameroonensis* (MLBs=100%) was resolved as very close sister with high support (MLbs=97%) to *R. brunneoannulata*.

The ITS phylogeny (Fig. 2) poorly resolved the relationships among the investigated species, but at species level all individual species received high support values, including the still undescribed R. sp1, sp2 and sp3. The singletons *R. cibaensis* and *R. afrovinacea* obtain their species recognition from the considerable branch length and the very high supports obtained in the sister terminal clades. *Russula radicans* formed a highly supported clade (MLBs 90%), but only its forma *sankarae* obtained support threshold (MLbs=71%) differing in one consistant base pair from both other forms (fo. *radicans* and fo. *miomboensis*).

**Fig. 2.**
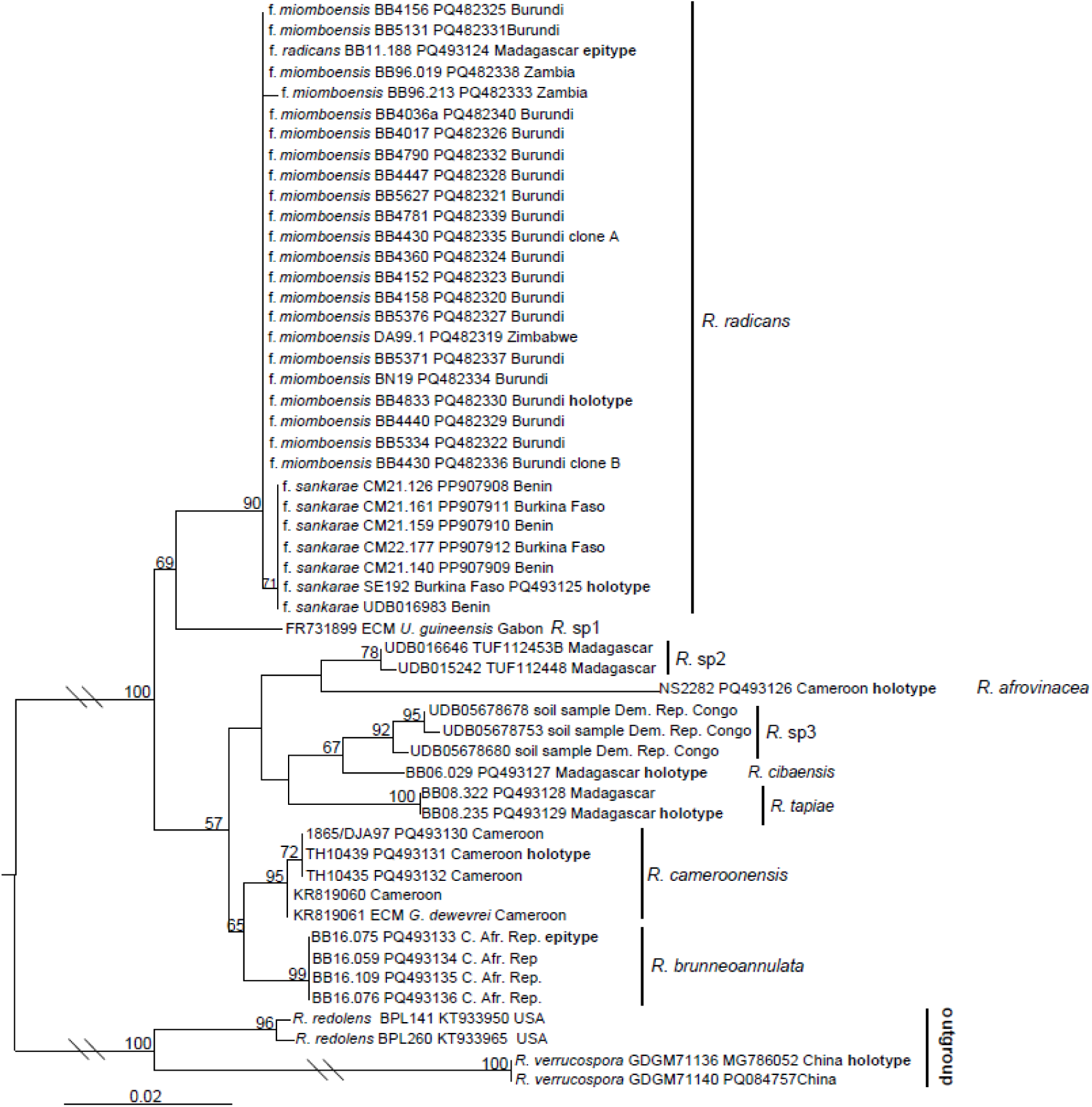
The best tree obtained by the Maximum Likelihood analysis of the ITS dataset for members of the new subsection, rooted with *Russula redolens* and *R.verrucospora* (two samples each). Bootstrap values ≥ 50% are indicated on the branches.

### Taxonomy

All examined species are presented below in alphabetical order. Subsections are discussed in the conclusion following the descriptive part.

***Russula acriannulata*** Buyck & Härkönen, *Karstenia* 33: 19 (1993) — Fig. 3

**Fig. 3.**
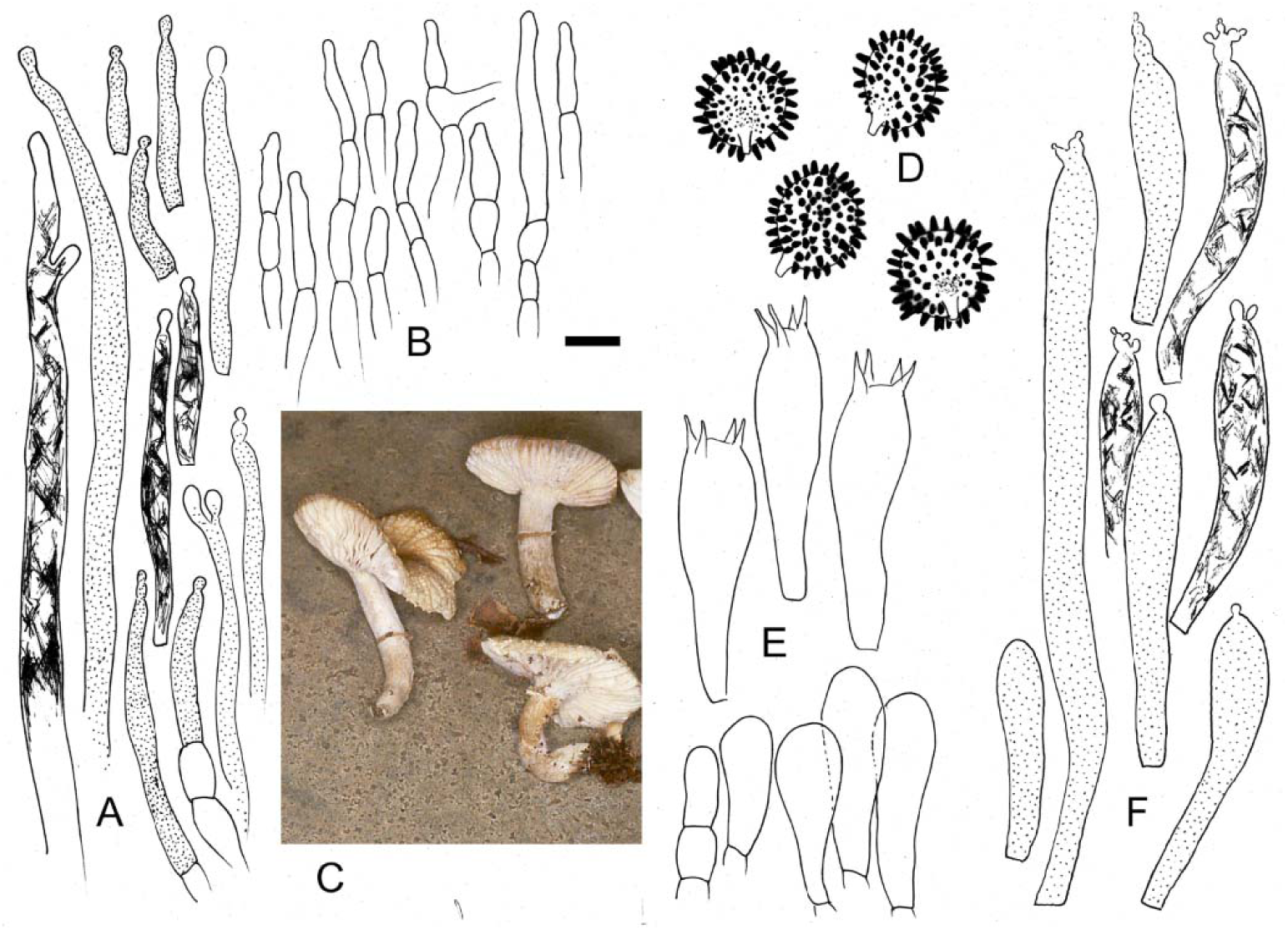
***Russula acriannulata* (**holotype). A. Long cystidioid hyphae from subpellis and shorter pileocystidia from near the pileus surface. B. Hyphal extremities near the pileus center. C. Fresh basidiomata. D. Basidiospore ornamentation. E. Basidia and basidiola. F. Pleurocystidia and a single cheilocystidium (left under). Scale bar = 1 cm for fruiting bodies, 10 µm for microscopic elements, but only 5 µm for basidiospores. Drawings: B. Buyck. Photo credit. M. Härkönen)

*Typus:* Tanzania. Tanga Prov.: Lushoto Distr., W. Usambara Mts, Mazumbai, N of the Mazumbai Forest Reserve border, N of the Kambi Falls (04 38 OC), lower montane forest, 1640 m asl, 8 Dec. l989, *Härkönen 10112* (H, isotypus PC0125177); ibid, 1640 m alt., T.Saarimaki 421 (paratypus H, isoparatypus PC0125178)

**Pileus** up to 40 mm in diam, ochre, darker brownish in the center; margin strongly tuberculate-striate. **Lamellae** white, up to 3 mm broad, with few lamellulae. **Stipe** annulate, white, tapering towards the base. **Flesh** very thin, not yellowing. **Taste** at first mild, becoming almost acrid. **Smell** unremarkable. **Spore print** not obtained (certainly pale).

**Basidiospores** subglobose to broadly ellipsoid, (8.4–)8.76–**9.14**–9.53(–9.9) × (7.4–)7.78–**8.11**–8.43(–8.7) µm (Q = (1.08–)**1.11**(–1.17)), densely omamented with very prominent, rather blunt, isolated verrucae, measuring up to 1.5 µm high, usually strongly amyloid, mixed with smaller verrucae; suprahilar spot indistinct, occupied by small, weakly amyloid verrucae, becoming larger towards the margin. **Basidia** voluminous, 35–55 × 12–16 µm, often rather short, four-spored. **Hymenial gloeocystidia** abundant, especially near the sterile gill edge, there 35–65 × 6–10 µm and often staining yellowish, on the sides of the gills longer (but difficult to measure because of their fragility), hardly emerging, clavate to subcylindrical, variously capitulate and usually with several smaller ‘buddings’ sprouting from the tip, thin-walled; contents well differentiated, abundant, typically giving the impression of being coiled against the inner wall leaving optically empty ‘windows’, sulfovanilline-positive. **Marginal cells** not observed. **Trama** of the gills composed entirely of sphaerocytes, but with long, cylindrical cystidioid hyphae. Subhymenium pseudoparenchymatic. **Pileipellis** very thin, subpellis hardly differentiated from the underlying trama, composed of hyphae with thin but weakly encrusted walls and with extremely abundant, prominent pileocystidia that become cystidioid hyphae in the underlying trama, these up to several hundreds of µm long, 6–9 µm in diam, broadly rounded to capitate, with distinct contents similar to hymenial cystidia; suprapellis a loose trichoderm composed of very fragile and thin-walled, short-celled hyphal extremitis, composed of easily collapsing cylindrical to ellipsoid or otherwise inflated cells, 3–5(8) µm diam.; pileocystidia in suprapellis very abundant, narrowly cylindrical to flask-shaped or even subulate, in the pileus center sometimes very short, 15–60(85) × 3–5(7) µm, capitulate, with distinct, similar contents as hymenial gloeocystidia, thin-walled. **Stipitipellis** very similar, caulocystidia equally abundant. **Clamp connections** absent.

*Notes*: The above description is borrowed from the original publication with some additional observations added on the basis of a new photograph and our microscopic re-examination. This East African species appears now morphologically quite different from *R. radicans* and allies by its (very) small size, white and practically equal lamellae, the thin flesh, the absence of any orange-yellow discolourations, the narrow and easily collapsing, short-celled hyphal terminations in the pileipellis, the even much more abundant gloeocystidia (both at the surface tissues and throughout the context) and, finally, the larger basidiospores with denser and higher spore ornamentation.

The newly obtained field photograph (Fig. 3C) much more clearly shows that the basidiomata are indeed not yellowing, but rather become grayish when handled or with age. Although it is still possible that *R. acriannulata* belongs to *Radicantinae*, its exclusion is also suggested by preliminary sequence data (of unsufficient quality) for mtSSU and LSU loci. When performing nBLAST on GenBank, both the nBLAST top scores and the minimum evolution tree placed this species near the American *R. redolens* Burl. (LSU) and the tropical Indian *R. shoreae* D. Chakr., A. Ghosh, K. Das & Buyck, (mtSSU), two closely related species near subsection *Substriatinae* X.H. Wang & Buyck (Gosh et al. 2023). One of our earlier multilocus ML analyses (not shown) included the obtained mtSSU and LSU sequences for *R. acriannulata* and placed this species near subsect. *Ilicinae* Buyck (a basal lineage in section *Heterophyllae*). The recently described *Russula indoilicis* Atalf et al. from Kashmir, India (Altaf et al. 2022) belongs in *Ilicinae* and shares an identical spore ornamentation with *R. acriannulata*. More high quality sequence data are still needed to place this African species with more accuracy.

**Russula afrovinacea** Buyck, T.W. Henkel & N. Siegel, **sp. nov.** — Figs. 4-5

**Fig. 4.**
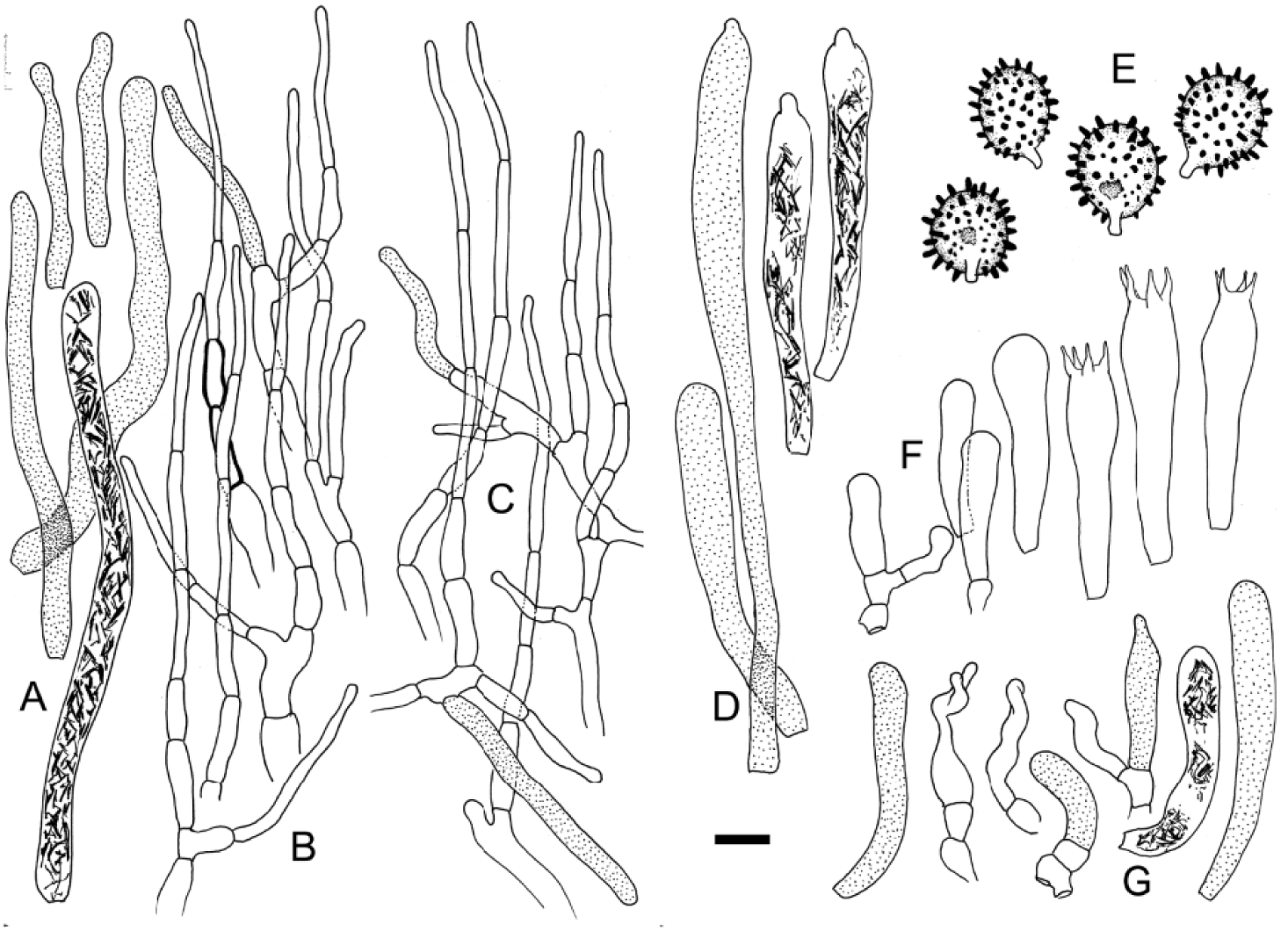
***Russula afrovinacea* (**holotype) A. Longer cystidioid hyphae from subpellis and shorter pileocystidia from near the pileus surface. B. Hyphal extremities near the pileus center. C. Hyphal extremities near the pileus margin. D. Pleurocystidia. E. Basidiospore ornamentation, notice also partially amyloid suprahilar spot. F. Basidia and basidiola. G. Cheilocystidia and marginal cells. Scale bar = 10 µm but only 5 µm for basidiospores. Drawings: B. Buyck.

**Fig. 5.**
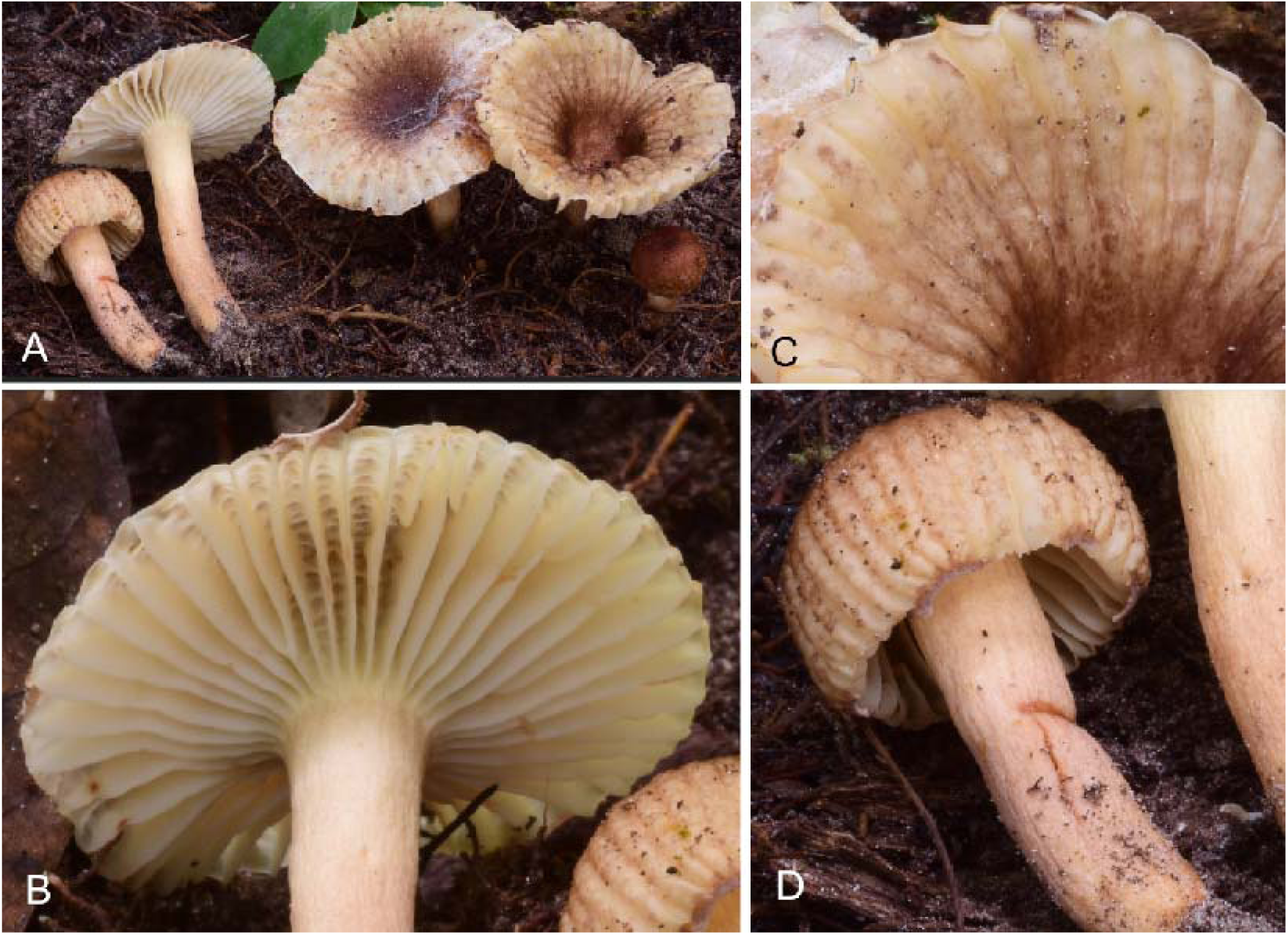
***Russula afrovinacea* (**holotype). A. Field aspect of the collection. B. Detail of the hymenophore showing the strongly concentrical veination between the lamellae. C. detail of the pileus surface at maturity; notice the yellow discoloration of the underying context near the margin. D. Young specimen with part of annulus attached. Photo credits: N. Siegel.

*Diagnosis:* differs from other species in the subsection by the combination of a predominantly reddish brown to brownish vinaceous colours of the pileus, the more regularly distributed high warts or spines on the spores, the very slender, multiseptate hyphal extremities of the pileipellis and its association with *Uapaca* species in humid forest.

*Holotype:* CAMEROON. Dja, 50 m east of base camp, scattered in sandy soil in riparian zone, under *Uapaca*, 11 Sept 2017, N. Siegel 2282 (YA **holotypus;** PC0125179, HSC-F 004445, isotypi)

*Etymology:* named after the African origin and vinaceous colour.

*Mycobank xxxxxxxxx*

**Basidiomata** rather small to medium-sized, fragile, growing dispersed in small groups. **Pileus** 26–51 mm diam., nearly globose and enclosing the stipe apex when young, later becoming depressed centrally and with uplifted margin; margin strongly tuberculate-sulcate, extremely thin-fleshed, carrying attached remnants of the annulus; pileus cuticle not peeling, faintly moist to dry, the center finely matted-tomentose and dark reddish brown (7F5–6), later somewhat fading (6E6–7); near the margin paler, orange brown (6D5) when young, then gradually becoming glabrous and cream coloured (4A3) as the pileus expands, staining orange on injury (5C6). **Annulus** membranous, concolourous with pileus center, breaking up in fragments that remain attached to the pileus margin, whitish, pruinose. **Lamellae** adnate or shortly decurrent with tooth, subdistant, subequal and intermixed with occasional lamellulae of variable length, some forking, concentrically and strongly interveined in the dorsal zone between lamellae already from the very young stages as evidenced in still closed caps, 5 mm high at mid-radius, not rounded near the pileus margin, cream (4A2–3) but staining orange-red (6B6–7) on injury; edges concolorous, even. **Stipe** 22–38 × 5–7 mm, cylindrical, under hand lens faintly pruinose from brown punctuation, cream (4A3), blushed with brownish (6D3) near base, bruising orange-red (6B6–7); stipe interior lacunar, soon hollowing. **Context** very thin, whitish to cream. **Taste** mild. **Smell** indistinct. **Spore deposit** white.

**Basidiospores** shortly ellipsoid, (7.71)8.03–**8.48**–8.93(9.38) × (6.04)6.62–**7.01**– 7.40(7.71) µm, Q = (1.11)1.15–**1.21**–1.27(1.30), ornamentation composed of isolated, conical to cylindrical spines and low warts, not particularly dense, up to 2 µm high; suprahilar spot in its distal part frequently weakly to moderately amyloid. **Basidia** mostly 41–50 × 9–11 µm, slender, clavulate, four-spored. **Marginal cells** mixed with cheilocystidia, slender, tortuous or undulate and less voluminous than basidiola. **Hymenial gloeocystidia** rather abundant, 1500– 2000/mm2, on the lamellae sides of very variable size, the longest ones deeply imbedded in the trama and > 100 µm long, most others measuring 51–77 × 9–10 µm, on the lamella edge also much smaller and measuring sometimes merely 25 × 6 µm; all cystidia thin-walled, clavulate, obtuse-rounded or papillate at the tip, filled with abundant, coarsely crystalline, SV-positive contents. **Subhymenium** small-celled, pseudoparenchymatous. **Pileipellis** obscurely two-layered, composed of a shallow, ill-developed suprapellis of intensively ramifying, very slender, ascending hyphal terminations and a thin subcutis of horizontal hyphae intermixed with many, very conspicuous cystidioid hyphae. A well-differentiated pseudoparenchyma (as in *R. virescens* for ex.) is not formed at the basis of the suprapellis. Hyphal terminations composed of one to six, predominantly thin-walled and narrowly cylindric cells; the terminal cell – usually also the longest one – measures 25–40(50) × 2–3 µm; cells beneath it are becoming slightly wider and shorter toward the basal cell, the latter frequently branching and mostly 5–7 µm wide. Pileocystidia measuring (20)40–100(150) × 5–8 µm in the pileus center, somewhat narrower at 3–5(8) µm wide toward the pileus margin, obtuse-rounded at the tip, never mucronate, often somewhat sinuous in outline, the smallest ones near the surface and terminal on hyphal extremities of the suprapellis, the longer ones originating from below the suprapellis, often remaining submerged as cystidioid hyphae in the subpellis and underlying pileal trama and there often several 100 µm long, cylindrical, not branching, filled with abundant, coarsely crystalline, SV-positive contents. **Clamp connections** absent.

*Notes:* This new species shares the typical attributes of the subsection with the other species, although lamellulae and forkings are less frequent compared to some other species (e.g. *R. radicans* or *R. brunneoannulata*). It is distinctly annulate, although the annulus disrupts always in larger fragments clinging to the pileus margin, and it shares the higher and more regularly dispersed warts on the basidiospores with the other African rain forest members of the subsection. Although the field notes mentioned an orange-reddish discolouration when bruised, this discoloration seems less intense than in some of the other members of this species complex.

*Russula afrovinacea* is unique in the subsection because of the predominantly reddish brown to brownish vinaceous pileus colour. The long and narrow hyphal extremities of the pileipellis are reminiscent of those of Malagasy *R. cibaensis* sp. nov. (with strong support [MLbs=93%] its closest relative in our multigene phylogeny, Fig. 1). To a lesser degree its hyphal extremities resemble also those of *R. tapiae* sp. nov., again from Madagascar. However, *R. afrovinacea* and *R. tapiae* have pileocystidia that are obtuse-rounded at the tip, while in *R. cibaensis* these are mostly capitate-mucronate, a difference that needs confirmation from future collections for these species. The ITS phylogeny (Fig. 2) reveals the existence of two related, but still undescribed species (R. sp2 and R. sp3) represented by environmental sequences. Finally, the *Uapaca* host habitat might indicate host specificity as in the case of some other *Radicantinae* or the unrelated *R. xylophila* – see discussion).

***Russula brunneoannulata*** Buyck, *Bull. Jard. Bot. Nat. Belg.* 60: 208 (1990). — Fig. 6-7

**Fig. 6.**
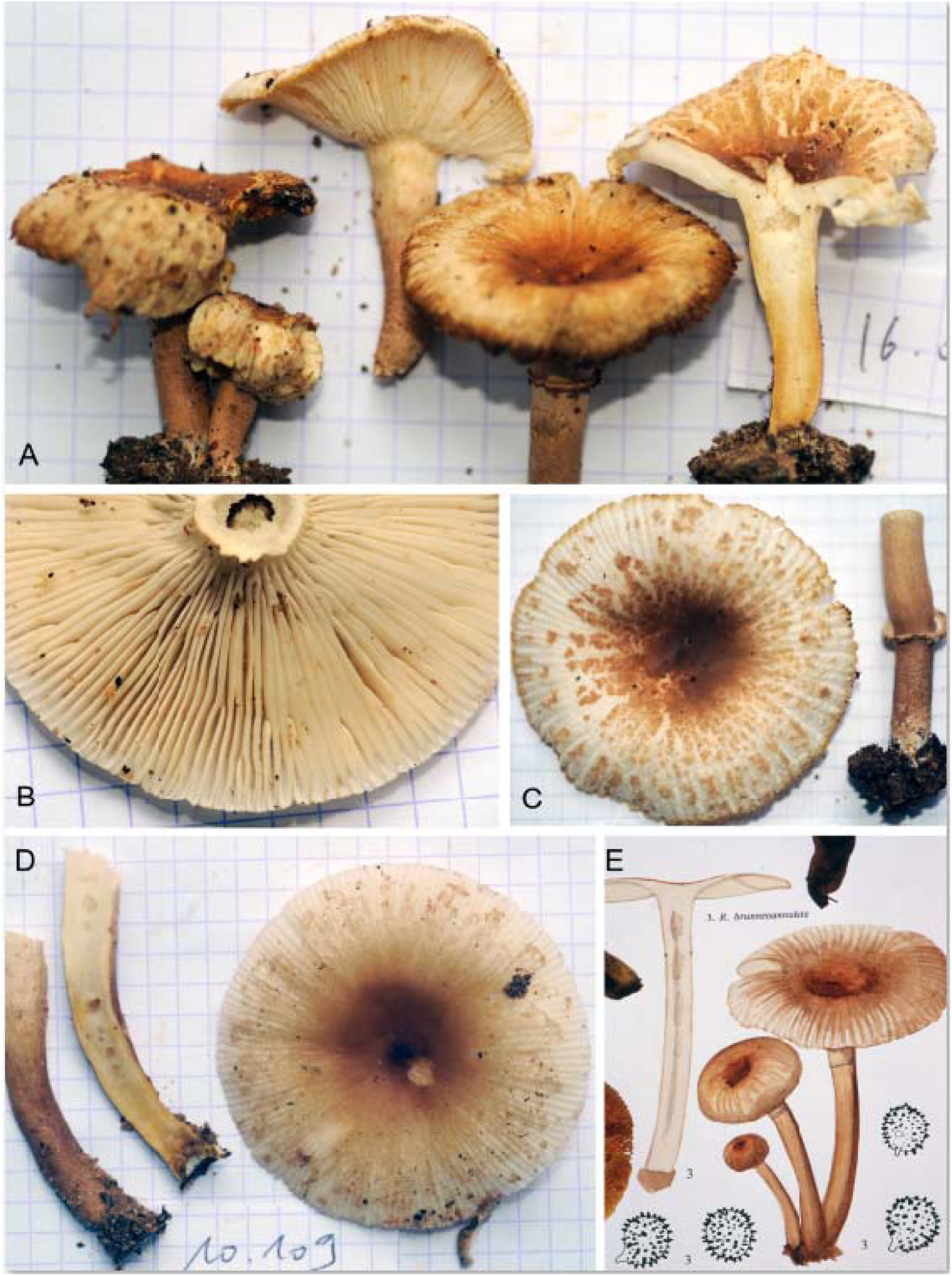
***Russula brunneoannulata***. A. Holotype specimens. B. Detail of the hymenophore showing the presence of many lamellulae and forkings in a rather dense arrangement compared to some of the other species. C. Pileus surface and stipe (BB16.076). D. Pileus surface with smoother aspect of the when more humid and sectioned stipe showing context discolouration (BB16.109). E. Original watercolour by Mrs. Goossens-Fontana and spore ornamentation, both from the holotype collection (from Buyck 1994, courtesy Bot. Gard. Meise). Photo credits: B. Buyck

**Fig. 7.**
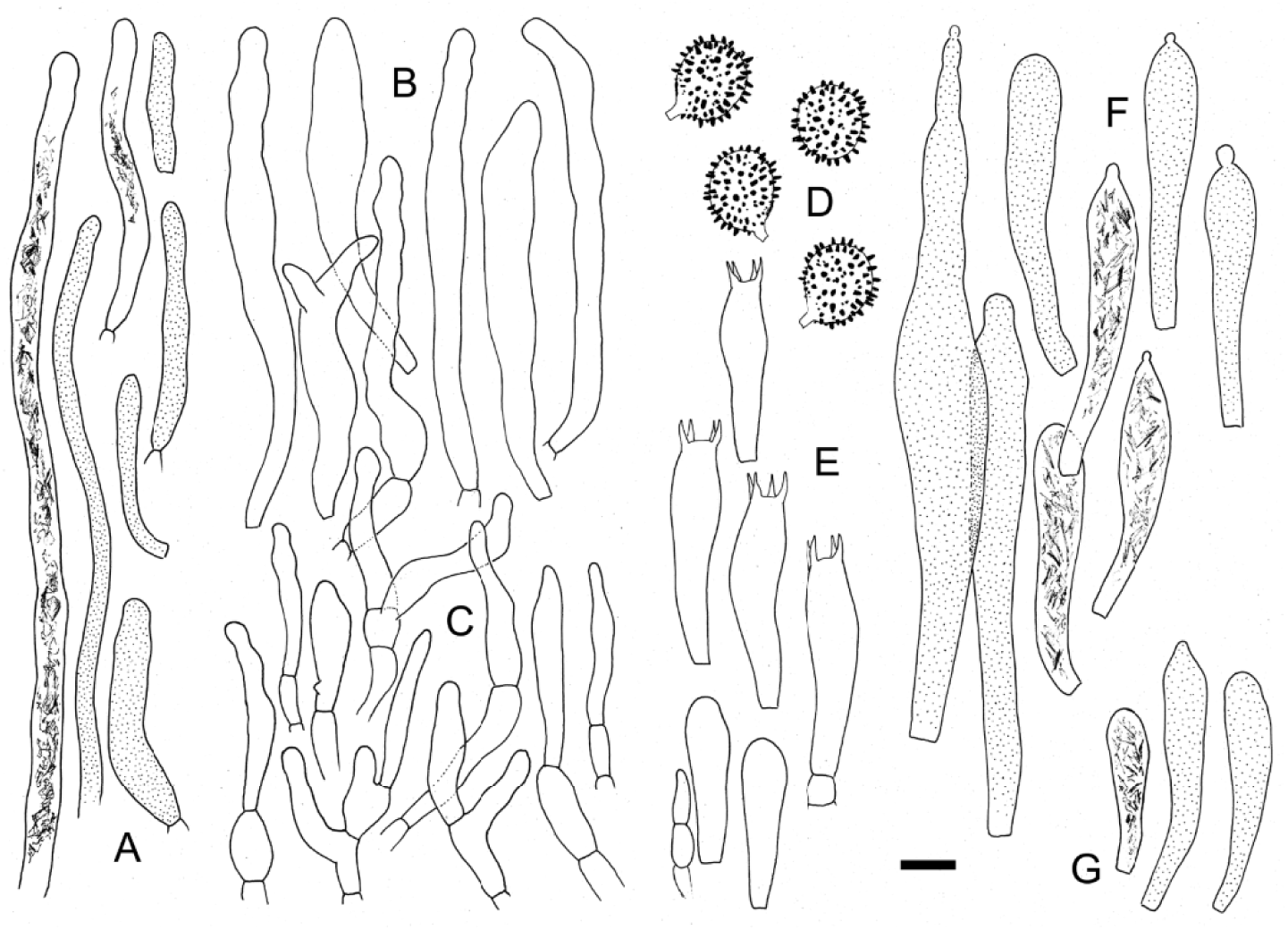
***Russula brunneoannulata*** (epitype). A. Long cystidioid hyphae from subpellis and shorter pileocystidia from near the pileus surface. B. Terminal cells near the pileus margin (above) and center (below). C. Spore ornamentation. D. Basidia and basidiola. E. Pleurocystidia. E. Cheilocystidia. Scale bar = 10 µm but only 5 µm for basidiospores. Drawings: B. Buyck.

*Original diagnosis*: *a R. oleifera Buyck differt carpophoris fragilioribus haud carnosis annulatis, pileipelle squamis pelliculisque destituta, cystidiis distincte latioribus, sporis elementis sejunctis cylindratis vel conicis ornatis, 7, 78 ± 0,61 × 6,88 ± 0,68 µm (Q=l,13)*.

*Holotype*: *Democratic Republic of the Congo: Binga, in groups on the forest soil or on wood in the Guineo-congolian rain forest, January 1935, Goossens-Fontana 1005 (holotypus BR)*.

*Epitype*: Central African Republic, near base camp Bai Hakou, Dzanga-Sanga national park, on sandy soil under *Gilbertiodendron dewevrei*, 19 May 2016, Buyck 16.075 (PC0125181)

MBTx xxxxxxx

*Description of the epitype:* **Pileus** 35–78 mm diam., center dark chocolate brown to rust brown, reddish brown, cinnamon brown (6CD4–7), rapidly much paler, yellowish brown to ochre or pale yellow and breaking up in very thin, appressed scales toward margin, exposing the whitish tissue below, strongly tuberculate-striate over 1/3 to ½ radius. **Lamellae** adnate, irregularly unequal with lamellulae in almost equal numbers, forkings present especially closer to stipe, not very dense (mostly less than 8 L+l/mm when fully mature), concentrically veined in the interstitial dorsal parts, ivory white to pale cream, rapidly discolouring yellowish when injured; edge concolourous, even. **Stipe** 36–65 × 6–9 mm, subcylindrical or slightly tapering downward, yellowing in lower part, concolourous with the pileus over most of its surface (6CD4–6), paler close to the lamellae and white at the extreme base, annulate, multi-chambered within, hollowing. **Context** very thin in pileus, firmer in the stipe, white, yellowing on injury. **Odour** none. **Taste** mild. **Spore print** white.

S**pores** 7.29–**7.53**–7.77(8.13) × (5.83)5.94–**6.15**–6.35(6.46) µm, Q = (1.13)1.18–**1.23-**1.28(1.32), ornamented with isolated small, pustulose warts and longer, rod-like to conical elements up to 1(1.5) µm high, strongly amyloid; suprahilar spot not or distally amyloid, often verruculose. **Basidia** mostly 32–43 × (8)9–10 µm, rather slender, clavulate, four-spored; sterigmata rather small, ca. 3–4(5) × 1–1.5 µm. **Hymenial gloeocystidia** numerous, emergent up to 50 µm, very different in size, near the gill edge originating in the subhymenium and small, sometimes only 35 × 8 µm, often clavate with obtuse-rounded tip; on the gill sides frequently continuing deep in the lamellar trama, mostly 60–150 × 7–12(16) µm. clavate to subfusiform-pedicellate, often appendiculate, or mucronate thin-walled; contents granular-oily, yellowish, reacting quite strongly to sulfovanillin. **Marginal cells** not observed.

**Pileipellis** orthochromatic in cresyl blue; subpellis poorly gelatinized, composed of narrow hyphae 2–3(4) µm wide and intermixed with abundant ascending pileocystidia up to 4 µm wide and immerged larger cystidioid hyphae (up to 8 µm wide) that continue in the pileus trama underneath; suprapellis becoming discontinuous toward the pileus margin exposing the subpellis with its many submerged pileocystidia and cystidioid hyphae; in the pileus center forming a dense mat of short, sometimes branched chains of 2–3 thin-walled cells; terminal ones 25–58 × (4)5–10 µm, subcylindrical to fusiform, often undulate in outline, mostly longer but not particularly wider than subterminal cells; toward the pileus margin with larger terminal cells, many between (50) 60–80(110) µm long and up to 13 µm wide, with occasional diverticulae. Pileocystidia thin-walled, without septa, near the pileus surface widely dispersed, but easily recognized by their strongly refringent contents, those originating as terminal cells in the suprapellis frequently very short, e.g. 25–30 × 3–5 µm, cylindrical, obtuse rounded at the apex, those that ascend from the subpellis equally wide, but rapidly much, much longer, up to several 100 µm, those near or in the pileus trama up to 8 (10) µm diam.; contents finely crystalline-granular, yellowish, SV-positive. **Clamp connections** absent.

*Additional examined material*. Central African Republic: near base camp Bai Hakou, Dzanga-Sanga national park, on sandy soil under *Gilbertiodendron dewevrei*, 18 May 2016, Buyck 16.059 (PC0714872); ibidem, 19 May 2016, Buyck 16.076 (PC0125182); ibidem, 23 May 2016, Buyck 16.109 (PC0125183).

*Notes*: Efforts to get at least a partial ITS sequence out of the holotype were unsuccessful. *Russula brunneoannulata* is genetically (98.5% similarity in ITS sequences) as well as morphologically very close to *R. cameroonensis* sp. nov. (see below) and both occur in the Central African rain forest. We hesitated for a long time whether to consider these as two different varieties of a single species, but it was impossible to get bootstrap support for this in our ITS phylogeny (Fig. 2). As far as field characters are concerned, it was important for our identification and epitypification that the original description of *R. brunneoannulata* did not mention any pinkish tinges and that the stipe was described as being of the same colour as the pileus. Field notes on the holotype further mentioned “relatively dense” lamellae, whereas the typical yellow-orange discolouration for *Radicantinae* was not even mentioned in the holotype field notes. Our new collections clearly confirm the close spacing of the lamellae compared with other *Radicantinae*, while the yellow-orange discoloration becomes only very obvious when sectioning the stipe, but remains overall slow and weak when handling the specimens. Compared to *R. cameroonensis* (see below) spindle-like cells are rare in the pileipellis of *R. brunneoannulata* as was already evident from the illustrations provided for the holotype (Buyck 1994, fig 268 where fusiform cells were not even represented) and are equally rare in the specimens studied here (Fig. 7).

***Russula cameroonensis*** Buyck & T.W. Henkel, sp. nov. — Figs. 8-9

**Fig. 8.**
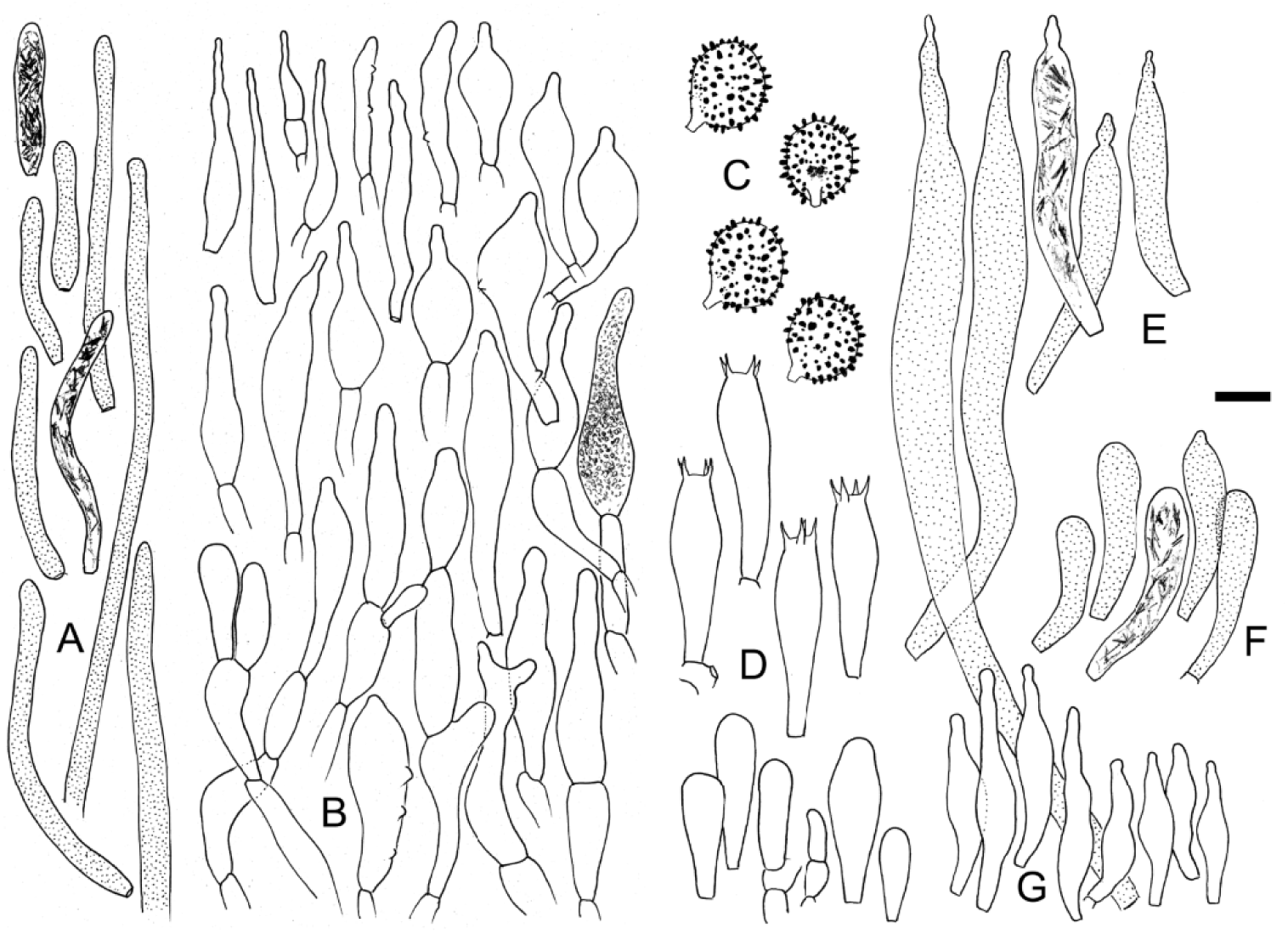
***Russula cameroonensis*** (holotype). A. Long cystidioid hyphae from subpellis and shorter pileocystidia from near the pileus surface. B. Terminal cells near the pileus center (upper half) and margin (lower half) with indication of pigment precipitation in one cell. C. Spore ornamentation. D. Basidia and basidiola. E. Pleurocystidia. F. Cheilocystidia. G. Marginal cells. Scale bar = 10 µm but only 5 µm for basidiospores. Drawings: B. Buyck.

**Fig. 9.**
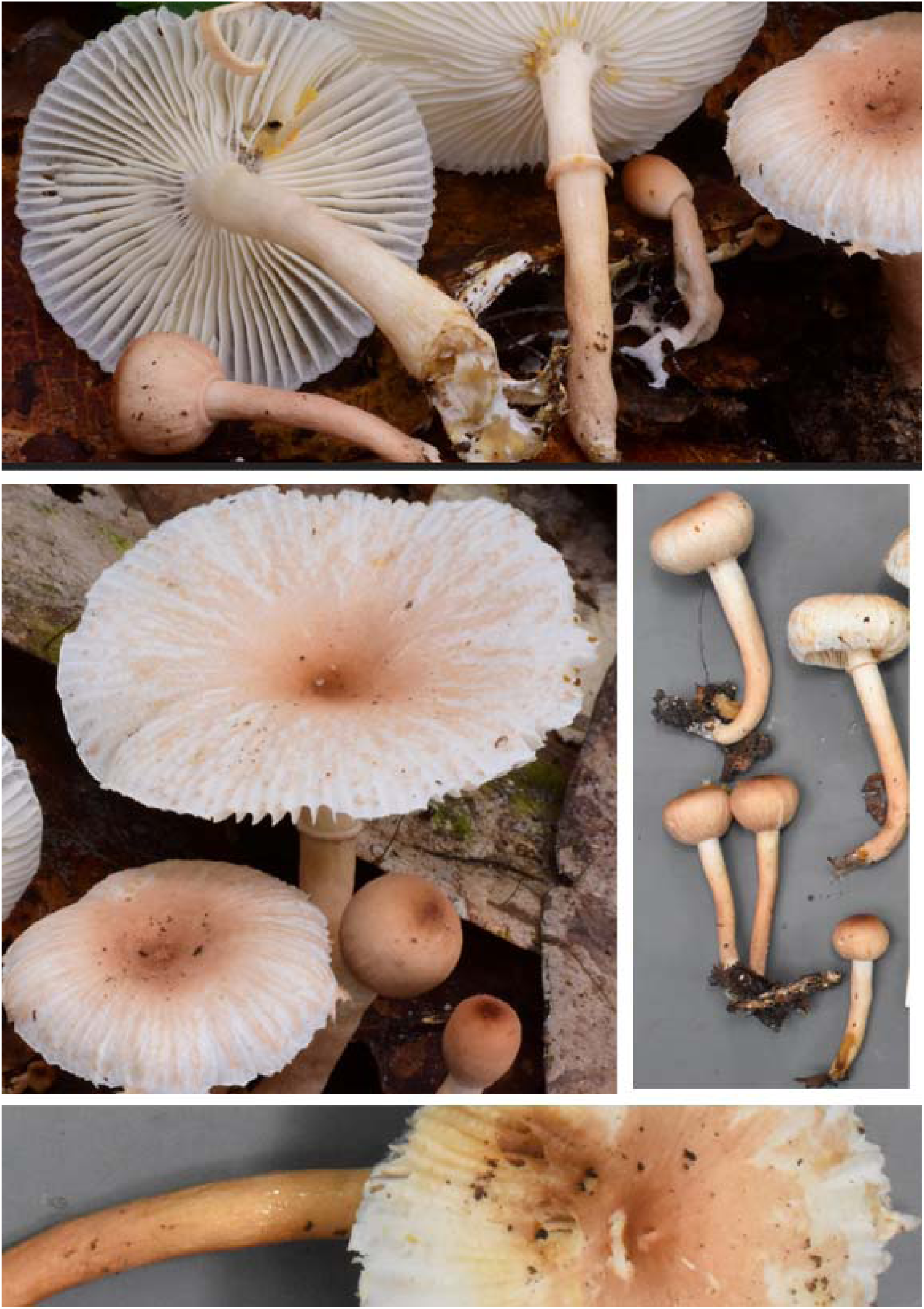
***R. cameroonensis.*** A-B. Field habit of the holotype collection, notice the unequal lamellae. C. Young fruiting bodies showing the flattened pileus center just before total expansion of the pileus (KM161) D. Close-up of fruiting body (KM161). Photo credits N. Siegel (A-B). T.W. Henkel (C_D).

*Diagnosis*: differs genetically from *R. brunneoannulata*, its closest relative and morphologically very similar species, by eight consistant base pairs (or. 98.6% similarity) in the ITS sequences. In the field it differs in the paler surfaces of pileus and lower stipe, which generally have distinct salmon or pinkish tinges; less densely spaced lamellae and a much more intense yellow-orange discolouration upon injury; microscopically, it differs in the predominantly spindle-shaped and overall smaller terminal cells in the pileipellis and similarly shaped marginal cells on the lamellae edges.

*Holotype: Cameroon.* Dja Biosphere Reserve, 2 km SW of base camp located at 3°21’29.8” N 12°43’46.9” W, in vicinity of *Gilbertiodendron dewevrei* plot 3, 29 Aug 2017, *T.W. Henkel 10439* (YA holotype; isotypes PC0125185, HSC-F 004446)

*Etymology*: ‘cameroonensis’ refers to the country it is described from.

*Mycobank*: MBXXXXXX

**Basidiomata** growing scattered to gregarious on humic soil, sometimes over several square meters. **Pileus** very thin-fleshed, 25–45(60) mm diam., young convex but flattened over the apex, remaining closed and attached to the top of the stipe until the latter reaches almost its full length, than expanding to broadly plane-convex to plane with a broad, shallow central depression, pileus surface overall pale pinkish orange (6A2–3) over the outer halve of its surface, toward the center progressively darkening to dark reddish brown (8DF5–6), deeply sulcate – striate and tuberculate over outer halve and with disrupted-areolate pileipellis fragments over a white layer underneath, not peelable or only over the outer part when young, in the center smooth, but minutely tomentose under handlens. **Lamellae** thin, subdistant, narrowly adnexed, 5–6 mm high, attenuated to 1 mm at the tips, unequal with mostly a single, 3–11 mm long lamellula between every 2 lamellae, rare forkings present, off-white to pale cream (4A2), discoulouring slowly orangish where bruised; edges concolorous, even. **Stipe** subcylindrical, long and slender, 27–60 × 3–7 mm, sometimes somewhat broader near apex (to 10 mm) and distinctly narrowed at the base (2–4 mm) which often shows white mycelial tomentum, over the lower 2/3 (under the annulus) light greyish orange (6A2–3, 5B3–4) with distinct shades of salmon pink and yellow, off-white above, glabrous and smooth, but under a handlens finely longitudinally striate, subsolid, stuffed - pocketed inside to distinctly chambered or hollow. **Annulus** inner side off-white, outer side concolourous with pileus, mostly remaining attached to the stipe, but sometimes disrupted in few fragments that remain attached to the extreme pileus margin, becoming free. **Context** off-white, yellowing with age or where injured, < 0.5 mm thick at pileus margin, 3 mm above the stipe. **Odour** faintly fragrant, sweet. **Taste** mild, pleasant. **Spore print** white.

**Basidiospores** subglobose to shortly ellipsoid, (6.67)6.88–**7.21**–7.53(7.92) × (5,42)5.70–**5.99**–6.28(6.46) µm, Q = (1.07)1.14–**1.21**–1.27(1.32), ornamented with isolated small, pustulose warts and longer, rod-like to conical elements up to 1(1.5) µm high, strongly amyloid; suprahilar spot not or distally amyloid, sometimes verruculose. **Basidia** mostly 33– 40 × (7)8–9 µm, rather slender, clavulate, four-spored; sterigmata rather small, ca. 3–4(5) × 1–1.5 µm. **Hymenial gloeocystidia** numerous,,emergent up to 50 µm, very different in size, near the gill edge originating in the subhymenium and small, sometimes only 35 × 8 µm, often clavate with obtuse-rounded tip; on the gill sides frequently continuing deep in the lamellar trama, mostly 60–150 × 7–12(16) µm. clavate to subfusiform-pedicellate, often appendiculate, or mucronate thin-walled; contents granular-oily, yellowish, reacting quite strongly to sulfovanillin. **Marginal cells** dispersed along the gill edge mixed with cystidia and basidia, slender and small, 24–42 × 5–7 µm, fusiformous, resembling the pileipellis terminal cells. **Pileipellis** orthochromatic in cresyl blue; subpellis poorly gelatinized, composed of narrow hyphae 2–3(4) µm wide and intermixed with abundant ascending pileocystidia up to 4 µm wide and immerged larger cystidioid hyphae (up to 8 µm wide) that continue in the pileus trama underneath; suprapellis in the pileus center forming a dense mat of short, sometimes branched chains of 2–3 thin-walled cells; the terminal cell 15–58 × 5–10 (13) µm, usually much more inflated compared to the subterminal ones and predominantly fusiform, lageniform, or ampullaceous, frequently somewhat constricted subapically (subcapitate) or abruptly narrowing into a longer beak, frequently also pedicellate, some irregularly sinuous-tortuous, mostly filled with some coarsely granular to drop-like contents (presumably pigment), becoming discontinuous toward the pileus margin exposing the subpellis with its many submerged pileocystidia and cystidioid hyphae. Pileocystidia thin-walled, without septa, near the pileus surface widely dispersed, but easily recognized by their strongly refringent contents, those originating as terminal cells in the suprapellis frequently very short, e.g. 25-30 × 3-5 µm, cylindrical, obtuse rounded at the apex, those that ascend from the subpellis equally wide, but rapidly much, much longer, up to several 100 µm, those near or in the pileus trama up to 8 (10) µm diam.; contents finely crystalline-granular, yellowish, SV-positive. **Clamp connections** absent.

*Additional examined material*: Cameroon. Dja Biosphere Reserve, within 2 km of base camp located at 3°21’29.8” N 12°43’46.9” W, under *Gilbertiodendron dewevrei*, 8 September 2014, leg. T.W. Henkel Dja97 (PC 0125186, HSC-F 004447); ibid., 28 August 2017, T.W. Henkel 10435 (PC0125184, HSC-F 004448), 23 September 2018, K.Mighell 161 (PC0125187, HSC-F 004449).

*Notes*: Although very similar in general aspect to *R. brunneoannulata*, this new species differs in the field already in the presence of distinct pinkish or salmon tinges on pileus and stipe, the overall more fragile, more slender and smoother aspect, the wider spacement of the lamellae and the much more intense yellow-orange discoloration when injured. Microscopically it is characterized by the predominantly fusiform terminal cells that are hyperabundant at the pileus surface and even distinctly present on the gill edge (Fig.8). Genetically, there is a consistent eight base pair difference in the ITS sequences between the two species. As for sequences obtained for the other loci, differences were limited to a single mutation in *tef1* and without consistent differences for the other genes.

***Russula cibaensis*** Buyck sp. nov. — Figs. 10-11

**Fig. 10.**
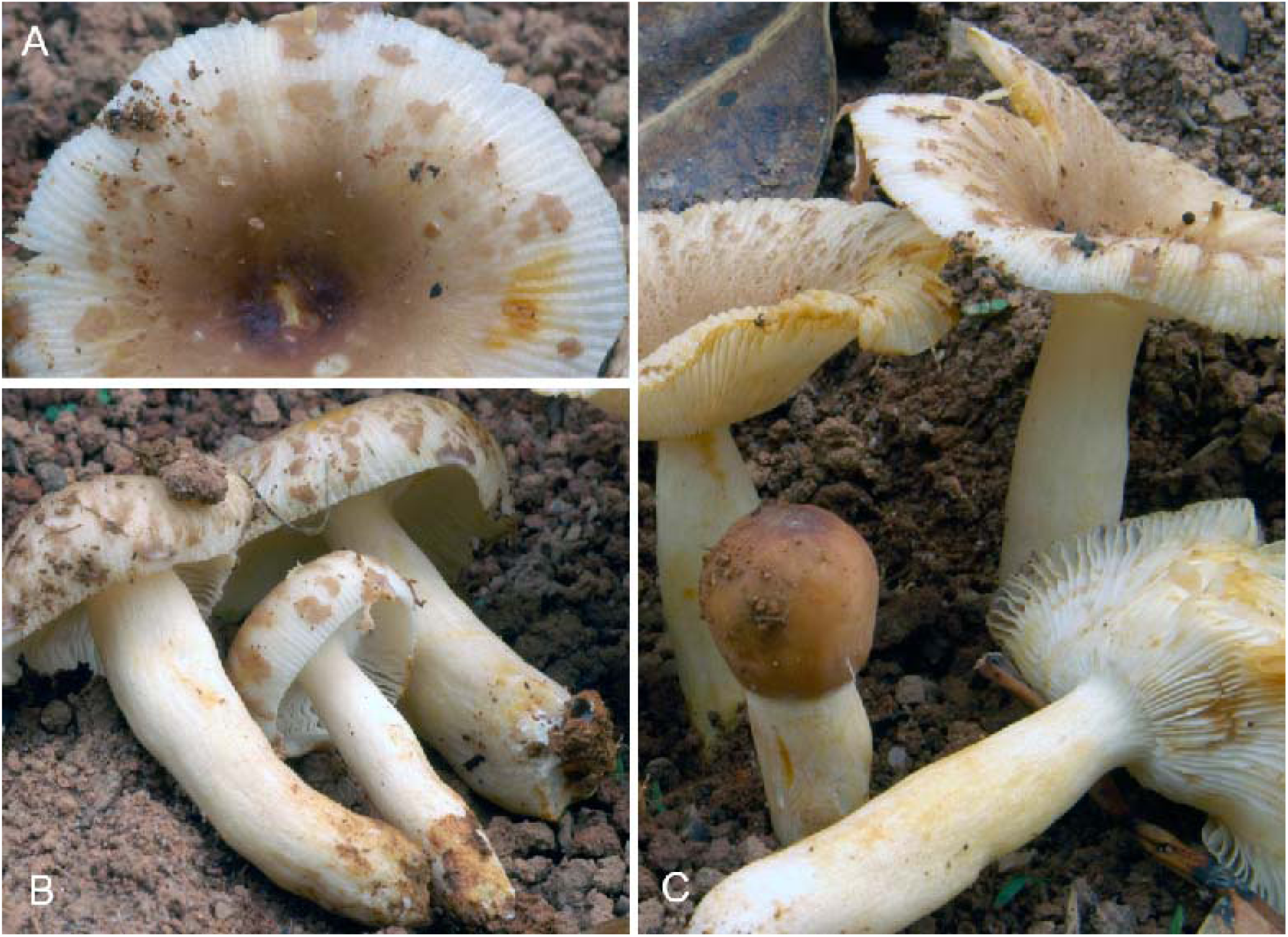
***Russula cibaensis* (**holotype). A. Detail of pileus surface; notice the energetic yellow-orange discoloration and the very thin and fragile pileus. B. Younger fruiting bodies during expansion with the annulus attached in fragments to the pileus margin. C. One young fruiting body and already strongly discolouring mature ones. Photo credits B. Buyck

**Fig. 11.**
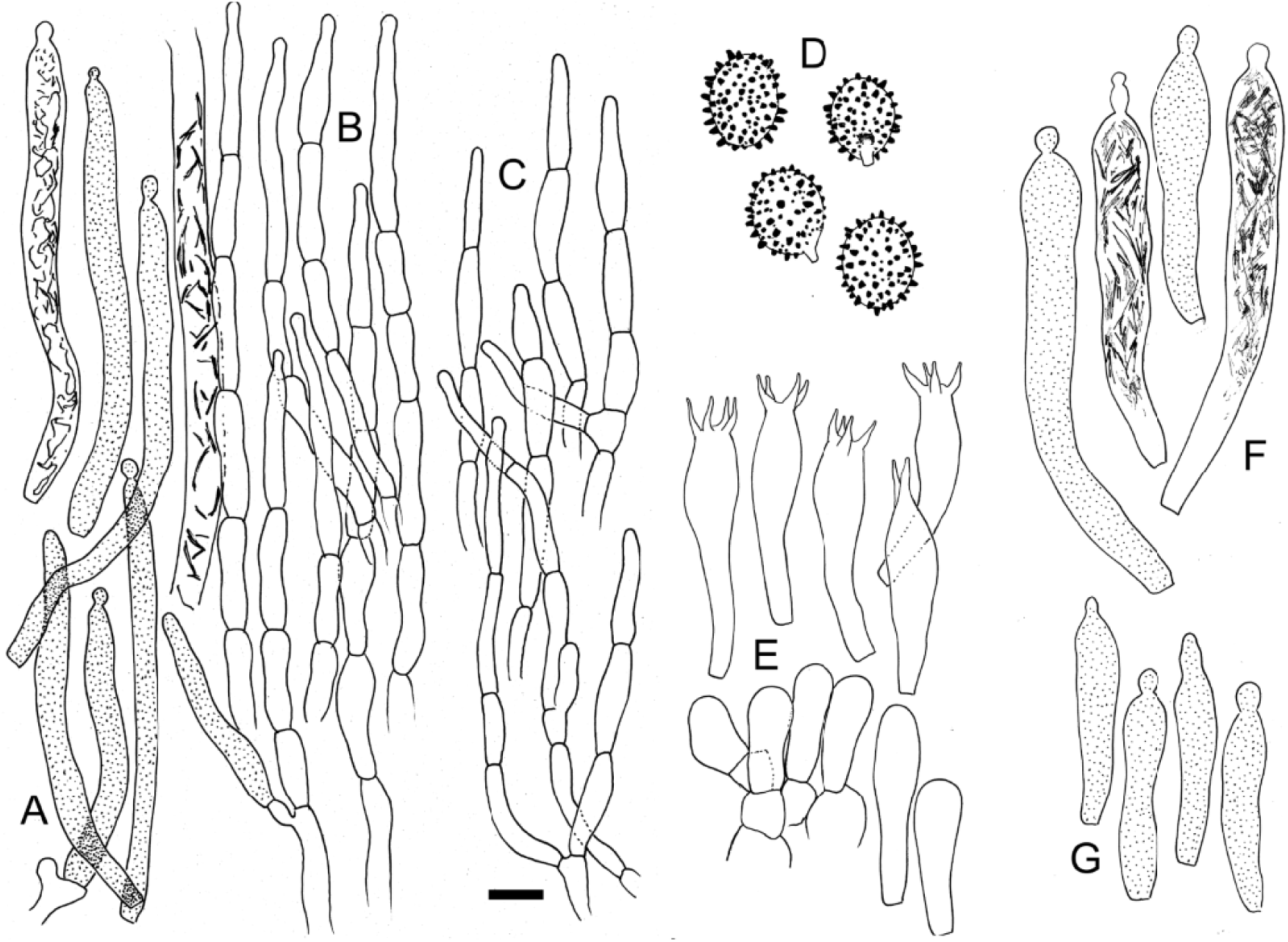
***Russula cibaensis*** (holotype). A. Pileocystidia of the pileus surface and a fragment of a cystidioid hypha from subpellis or underlying trama. B. Hyphal extremities of the pileus center. C. Hyphal extremities near pileus margin. D. Basidiospore ornamentation. E. Basidia and basidiola. F. Pleurocystidia. G. Cheilocystidia. Scale bar = 10 µm but only 5 µm for basidiospores. Drawings: B. Buyck.

*Diagnosis*. Differs morphologically from other *Radicantinae* in the humid forest biome by the combination of its distribution and habitat, the somewhat more pronounced acridity of its flesh and the unique brown tint of the pileus.

*Holotype*: Madagascar. Eastern escarpment, near Andasibe, on the old sawmill factory site (the “Complexe Industriel de Bois d’Andasibe” or C.I.B.A.), on bare ferralitic soil under *Eucalyptus robustus*, 21 January 2006, Buyck 06.029 (PC0124638, holotypus)

*Etymology*: named after the type locality near the old sawmill factory site (the “Complexe Industriel de Bois d’Andasibe” or C.I.B.A.),

*Mycobank*; XXXXXX

**Basidiomata** very fragile, growing in a group of more than ten dispersed specimens. **Pileus** 42–58 mm diam., rather irregular in outline, slightly depressed in the center, sometimes somewhat wavy outward and with azonate, strongly striate-tuberculate margin; surface glabrous but at the extreme margin distinctly pruinose under a handlens, dull, not hygrophanous, not viscose-glutinous, young separable except in the very pileus center, yellowish brown with dark chocolate brown disc when young, remaining darkest in the center, after expansion fragmenting rapidly toward the margin in a typical ‘virescens’ pattern, thereby exposing the whitish subpellis and context underneath. **Lamellae** normally spaced (ca. 1/mm), adnate to subdecurrent, 3 mm high, subequal from ‘more than usual’ lamellulae, with hardly any forkings except closer to the stipe, narrowing toward the pileus margin, with transverse veination in between the gills, white; edges concolorous, even. **Stipe** central, 50–54 × 9–12 mm diam., either somewhat shorter or longer than the pileus diam., subcylindrical to weakly conical or even ventricose, in its upper part frequently narrowed close to the gills; surface smooth, glabrous but with distinct pruina above the annulus (particularly when the pileus is still closed), white; the interior developing rapidly 2–6(10) cavities. **Annulus** well developed, brown on the lower surface, often remaining attached to the pileus margin after expansion. **Context** very thin, 2 mm thick above the gill attachment, white, yellowing when injured or upon handling, especially closer to the surfaces. **Taste** only moderately, sometimes tardily acrid. **Odour** insignificant. **Spore print** white or nearly so.

**Basidiospores** shortly ellipsoid, (7.71)7.95–**8.27**–8.59(9.17) × (6.46)6.81–**7.07**– 7.34(7.71) µm, Q = (1.11)1.12–**1.17**–1.22(1.29); ornamentation composed of isolated small, pustulose warts and longer, rod-like to conical elements up to 1(1.5) µm high, strongly amyloid; suprahilar spot not amyloid or distally so, verruculose. **Basidia** 36–45 × 8–11 µm, clavulate or most inflated in the median part, 4-spored, but particularly closer to the edge also frequently two and even one-spored. **Basidiola** slender, clavate. *Subhymenium* pseudoparenchymatic with relatively voluminous cells. **Hymenial gloeocystidia** numerous on lamella sides, 45–80(100) × (8)10–12 µm, near the lamella edge not exceeding 50 µm long, mostly with capitulate, rarely tapering-appendiculate tips, all thin-walled, with abundant and coarsely crystalline contents. **Marginal cells** not differentiated. **Pileipellis** two-layered, orthochromatic in cresyl blue; subpellis of intermingled, narrow hyphae mixed with numerous cylindrical, long cystidioid hyphae with abundant coarsely crystalline contents; suprapellis composed of slender, thin-walled, multi-celled hyphal extremities composed of 4–6(7) narrow, subcylindrical cells, mostly somewhat constricted in the median part, 4–6.5 µm diam.; the terminal cell (25)40–35–50 µm long, usually tapering upward and sometimes weakly but repeatedly constricted. Pileocystidia subcylindrical, long and narrow, (40) 45–90 × 5–7 µm, originating from branchings of subterminal cells or from ascending subpellis hyphae, obtuse or, more frequently, mucronate to minutely capitate at the apex, becoming cylindrical and very long in the subpellis to finally transforming into cystidioid hyphae in the pileus trama. **Clamp connections** absent.

*Notes:* Sequences for this species have been published previously (Buyck et al. 2018) when it was provisionally named ‘*R. cf brunneoannulata*’. The association with *Eucalyptus* remains to be verified, although it would not be so strange given the fact that this tree genus is in Madagascar associated with numerous (hundreds of) ectomycorrhizal fungi that made the host switch from endemic or African hosts to the Australian host (see Buyck 2008, Buyck in Aryiawansa et al. 2015), particularly so in the area where *R. cibaensis* was collected. Although eucalypts were the dominant trees at the collection site, we might have missed some young *Uapaca* or *Sarcolaena* saplings from the rain forest at very close distance. We are convinced that *R. cibaensis* is not an exotic (Australian) species that arrived in Madagascar together with eucalypts. This species has everything that is typical for some other *Radicantinae* from the humid forest biome: rather spaced lamellae with a strong concentrical veination in the dorsal zone, a very thin pileus context and energetic yellowing of the tissues upon injury and a spore ornamentation composed of rather spaced, medium high conical warts.

There are a few sequences obtained from soil samples in the eastern part of the Democratic Republic of the Congo that represent a closely related, still undescribed relative of *R. cibaensis* from the Central African rain forest biome (R. sp3 in Fig. 2).

***Russula radicans* forma *radicans*** R. Heim, *Candollea* 7: 389 (1938). — Figs. 12–14

**Fig. 12.**
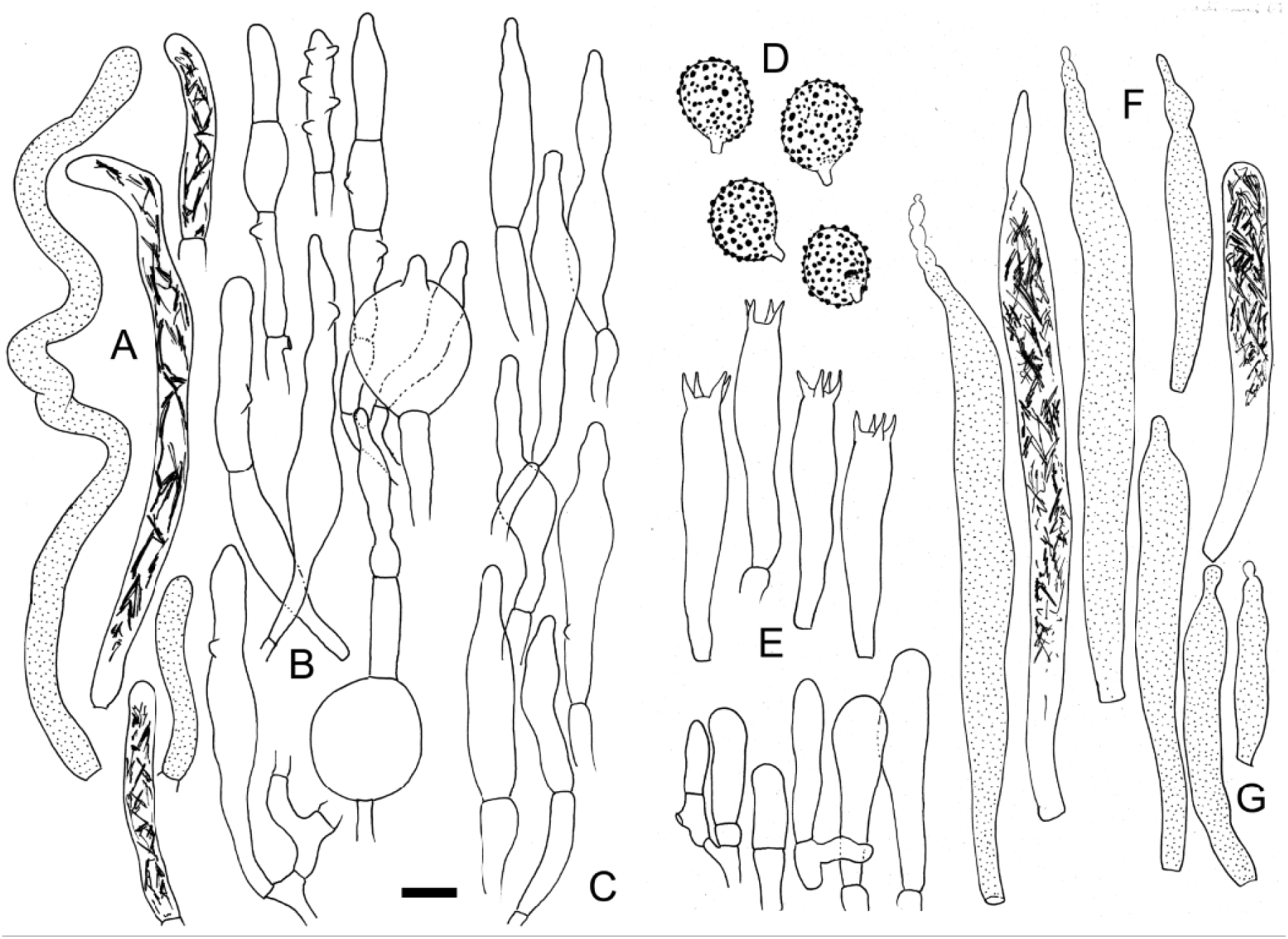
***Russula radicans*** (epitype). A. Longer cystidioid hyphae from subpellis and shorter pileocystidia of the pileus surface. B. Hyphal extremities of the pileus center. C. Hyphal extremities near pileus margin. D. Basidiospore ornamentation. E. Basidia and basidiola. F. Pleurocystidia. G. Two cheilocystidia. Scale bar = 10 µm, but only 5 µm for basidiospores. Drawings: B. Buyck.

**Fig. 13.**
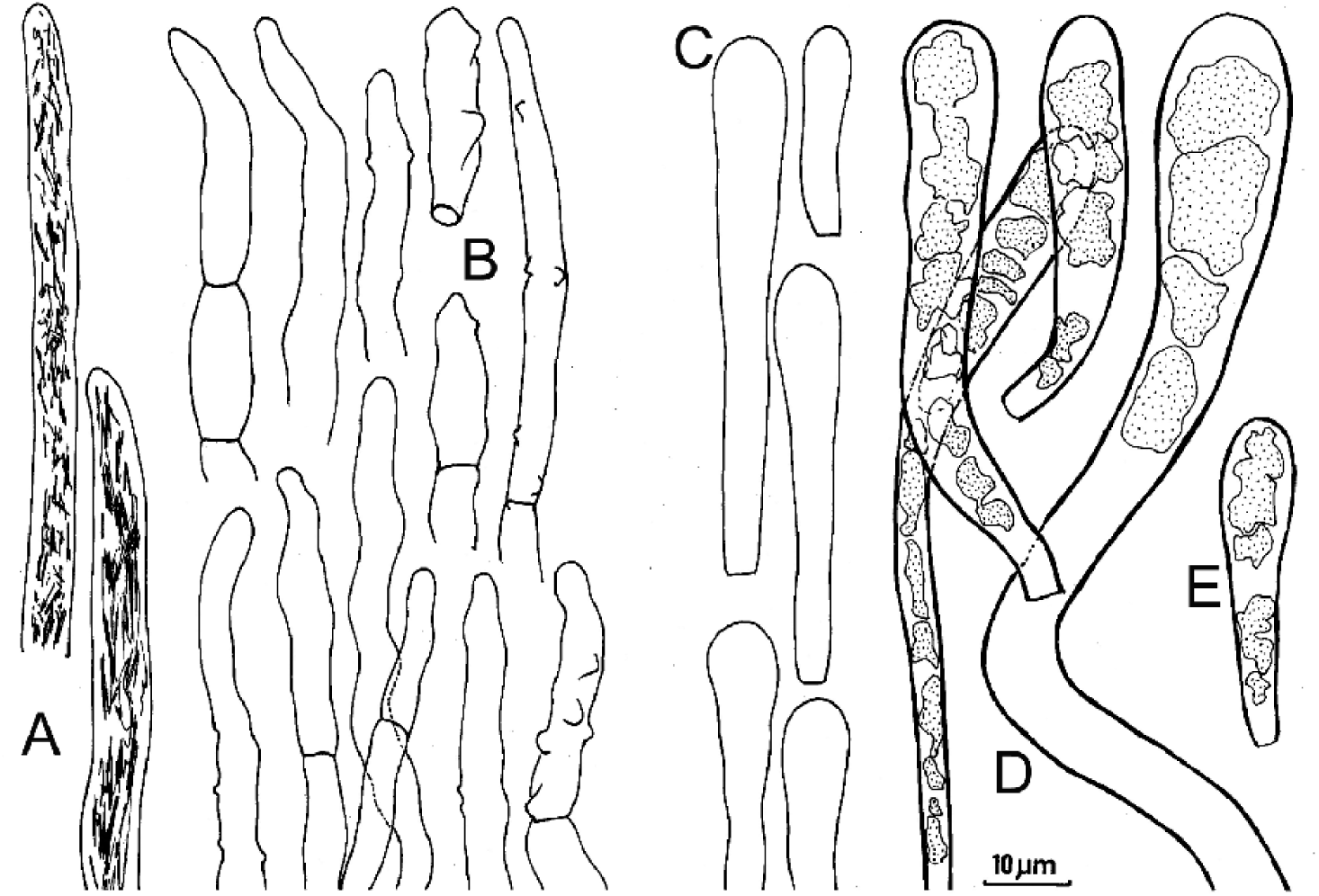
***Russula radicans*** forma *radicans* (holotype). A. Apical parts of cystidioid hyphae in subpellis. B. Terminal cells in pileipellis near pileus center. C. Basidiola. D. Pleurocystidia. E. Cheilocystidium. The poor state of the exsiccatum explains the absence of illustrations for basidia with sterigmata, as well as for hymenial cystidia with mucronate apices or strongly inflated cells in the pileipellis. Scale bar = 10 µm. Drawings: B. Buyck.

**Fig. 14.**
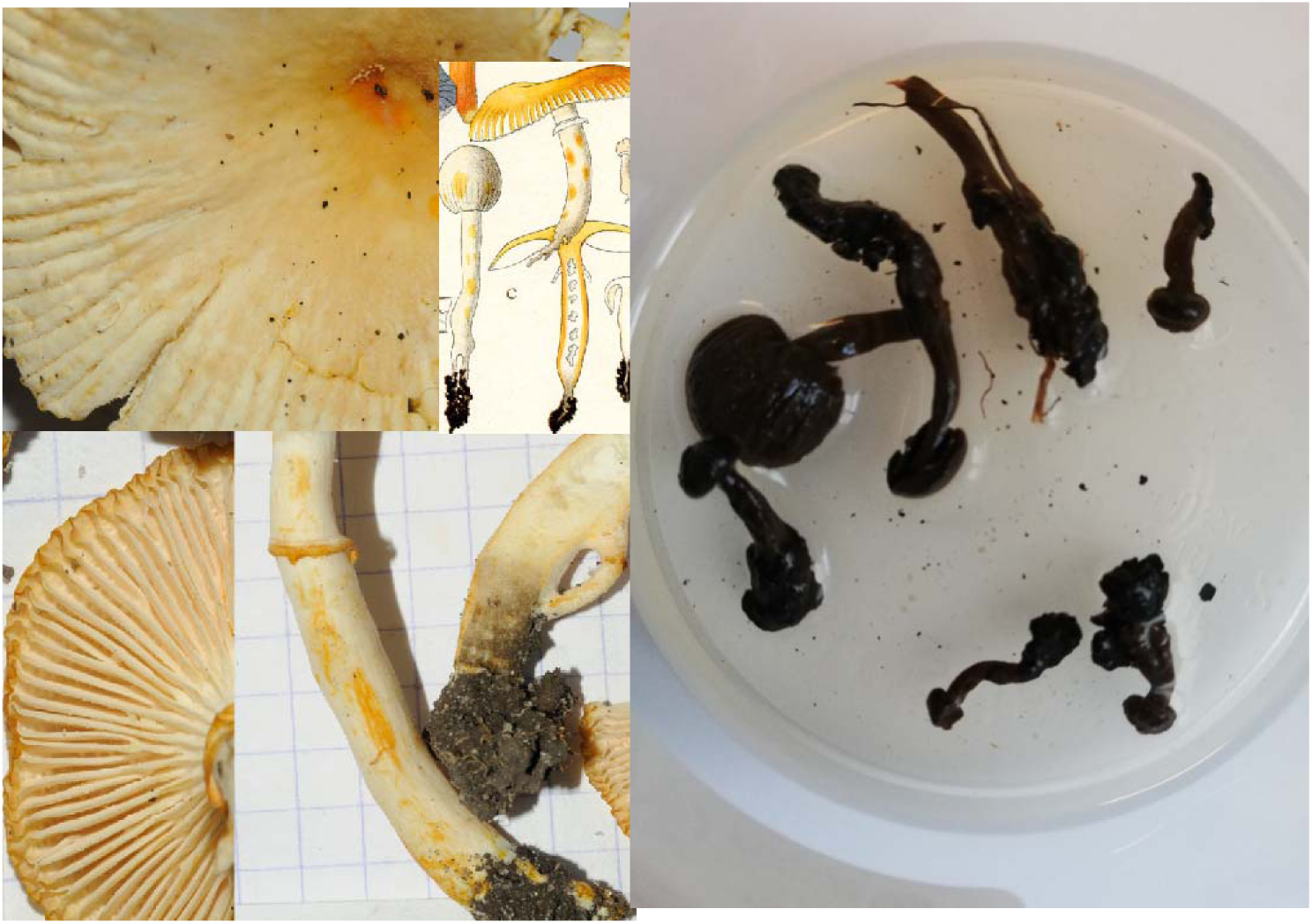
***Russula radicans*** (epitype & holotype). A. Detail of pileus surface. B. Watercolour of the holotype (taken from Heim 1937). C. Detail of lamellae showing the presence of furcations and lamellulae. D. Detail of stipe and annulus showing the intense orange-yellow discoloration on handling. E. The holotype, now again kept in conservation liquid (PC). Scale bar = 1 cm. Photo credits B. Buyck

*Original diagnosis*: ***Pileo*** *2–6 cent. diam., mox subpelliculari, primum tubuliformi subgloboso, dein valde convexo ; mox piano et subpelliculari, margine longe valde striata et subobtusa, velo generali albo gelatinoso caduco ; cute satis crassa, secernibili, subtiliter tomentosa; primum pallide aurantia, dein ex cremeo-roseo aurantia, mox ad marginem ex luteo aurantia, in medio intense aurantia, sub cute subrosea. **Stipite** in statu juvente 5-6 cent. alto, ad 6-7 mm. lato, ad basim inflato, longo, inaequali, radicato, subtiliter tomentoso, sursum striolato, primum ex cremeo roseo, in apice albo, dein roseo, aurantio colore tincto, in medio farcto-lacunoso.* ***Annulo*** *membranaceo, fere duplici, mobili, cerneo, adnato, non striato, superiore annulo descripto, aequali, parvo, inferior pendente, lacero, elato, permanenti vel caduco, interdum coronae descriptae modo circa marginem exstante. **Lamellis** parum confertis, parvis lamellulis singularibus intermixtis, subcrassis, angustis (∼ 3,5 mm), dente longo subadnatis, mox subliberis. **Carne** primum alba, dein ex luteo aurantia, tactu intense aurantia, brevi in cortice stipiti aurantia facta, deorsum nigricante, miti dein nauseoso sapore, ingrato non autem acri, inodora*.

***Sporis*** *in cumulo albo cremeis, breviter ovatis, 7,5-9,3 × 6,3-7,6* μ*, verrucis crassis humilibus distantibus obtusis (0,6-0,7* μ *alt.) singularibus ornatis. **Basidiis** 45-58 × 8, 7-9,5* μ*., elongatis-piriformibus, tetrasporis vel interdum bisporis. **Cystidiis** in acie et in lateribus praesentibus, 35-75 × 7, 5-13* μ*, cylindratis, fusiformibus vel longe piriformibus, in lateribus acutis vel sursum mucronatis, tegumento tenui et hyalino*.

*Ad terram humosam, gregatim vigens, radicans vel fasciculatus, in silva arenosa prisca littorali orientali insulae. Madagascar. Heim nr. G71, 7 décembre 1934*.

**Madagascar**. North of Fenerive, South of Soarienara, in sandy coastal forest, 7-9 Dec. 1934, Heim RH13 (=G71, PC0125178, holotype)

*Epitype*: Tampolo forest, ca 10 km N of Fénérive Est, South of Soanierana, in littoral forest, near the gard post at the entrance, in deep sandy soil under *Intsia bijuga*, 9 July 2011, Buyck 11.188 (PC0125159; **epitypus hic designatus**)

*Mycobank*: MBTxxxxxxx

*Description of the epitype*: **Basidiomata** fragile, instantly becoming intense yellow when handled or injured, growing in small groups. **Pileus** very thin, up to 43 mm diam., gently concave, margin strongly tuberculate-striate up to mid-radius, cuticle dull, under a hand lens powdery-farinaceous particularly in the center, there vivid orange-yellow, but rapidly fading to pale ochraceous or pale lemon yellow, fissuring outside the centre, but not forming well-recognizeable scales, cream or almost white near the margin. **Stipe** slender, 51–59 × 4–9 mm, subcylindrical, whitish to pale cream, the interior lacunar and hollowing with age. **Annulus** remaining mostly around the stipe or, more rarely, attached to the pileus margin. **Lamellae** adnate, normally spaced (ca 1/mm), subequal and irregularly intermixed with smaller lamellulae, furcations present here and there, ca 4 mm high, not rounded at the pileus margin, with interstitial concentrical veins in between lamellae; lamella edges concolourous, even. **Context** very thin and brittle, white, instantly becoming intense yellow to orange yellow when handled or injured. **Taste** mild. **Odour** unremarkable. **Spore print** not obtained but forming white deposits on lamellar surfaces.

**Basidiospores** shortly ellipsoid, (7.08)7.28–**7.63**–7.97(8.33) × (5.63)5.86–**6.23**– 6.60(7.08) µm, Q = (1.12)1.17–**1.23**–1.28(1.30) ; ornamentation composed of isolated, convex to conical warts, not very densely disposed, strongly amyloid, of unequal size, the largest up to 1 µm high ; suprahilar plage verruculose, inamyloid to distally amyloid. **Basidia** slender, 39–51 × 8–9 µm, (2)4-spored. Basidiola clavate, up to 10 µm wide, but finally narrowing again in their upper part before forming sterigmata. **Subhymenium** pseudoparenchymatic. **Hymenial gloeocystidia** abundant, ca 3000/mm2, (34)40–90(150), probably even longer sometimes with the longest originating from the lamellar trama, often running parallel with the surface just underneath the subhymenium, clavate, fusiform, with (sub)capitate to moniliform apices, more rarely obtuse-rounded. **Marginal cells** not observed. **Pileipellis** composed of a trichodermal suprapellis and an ill-defined subpellis mixed throughout with abundant pileocystidia; suprapellis of ascending and more or less aggregated, and rarely branching chains of two to three or four cylindrical to variously inflated cells; these often ellipsoid or inflated near one of the septa, up to 10 µm wide, few cells strongly inflated to almost globose and up to 20(25) µm wide, especially near the base of the trichoderm; terminal cells (15)25 – 70 × (5)7–10(12) µm, exceptionally wider, mostly attenuated upward, often undulate in outline or repeatedly constricted, with frequent diverticules. Pileocystidia abundant, very prominent because of their strongly refringent, crystalline contents, cylindrical, 5–6 µm wide, those constituting terminal cells in the trichoderma rather short, 30– 70 µm long, when ascending from or submerged in the subpellis much, much longer, often coiled and turning into cystidioid hyphae in the trama underneath. **Clamp connections** absent.

*Additional examined material*. **Madagascar**. Marotandrano, escarpment East coast, NW of Fenerive, no date, R. Heim nr. 1 (PC0714873); forest of Ivolaina, near Tamatave, 15 sept 1937, leg. Arsene Botozanany, in herb. R. Heim nr.11. (PC0714874)

*Notes:* Both the holotype, collected nearly one century ago (in 1934) by R. Heim, and the epitype were collected in the sandy soils of the littoral forest north of Fénérive, South of Soanierana, on Madagascar’s East Coast, possibly at the very same location. The degree of resemblance between the holo- and epitype collection is indeed so remarkable that one wonders if they don’t represent the same population, both sharing the extremely vivid colours and the same artefact of a stipe making a short lateral loop near its radicating base (fig. 14D). It could be important that *Intsia bijuga* (Fabaceae subfam. Detarioideae) was the most likely host tree for the epitype, which could, together with the deep sandy soil (as on a typical beach at the sea shore), possibly explain the morphological differences with genetically identical collections from other habitats on the African mainland (see below).

Although Heim’s original diagnosis does not give any specific details on the structure and composition of the pileipellis, the hyphal terminations were briefly described by Heim in his monograph on the Russulaceae of Madagascar (Heim 1938) as “erect, cylindrical, 3.5–7 µm diam., septate, with cells being 25–40 µm long and having a nodulose-sinuous outline”. The nodulose aspect mentioned by Heim is now usually referred to as “diverticulate” in European *Russula* monographs (Romagnesi 1967, Sarnari 1998). In Europe, the presence of diverticula is considered to be quite useful for the identification of some *Russula* species (such as for *R. cuprea* and close allies in subsect. *Urentes* Mre., an otherwise completely unrelated group of acrid species in the crown clade of *Russula* subgen. *Russula* - see Buyck et al. 2018). These diverticula were equally very conspicuous in the epi- and holotype of *R. radicans*, but much less so in the majority of the other *Radicantinae*, although they are nearly always present.

The holotype tissues (as those of the other collections made by Heim and collaborators for this species) barely inflated as these specimens had been stored in conservation liquid which completely evaporated over the years, resulting in brick-hard, black basidiomata (see fig. 14E, which illustrates the type presently again conserved in liquid). As a result, microscopic observation of holotype tissues has become extremely difficult, particularly for the very poorly inflating hymenium, whereas spore ornamentation became insensitive to Melzer’s reagent. We could, however, observe that pleurocystidia in the holotype were of similar variable size as in the epitype, i.e. often much longer than mentioned in Heim’s description. Heim’s description did not mention the ampullaceous cells dispersed here and there in the lower part of the suprapellis, but our examination of the holotype clearly confirms their presence, although these large cells hardly inflated and are, therefore, not represented in our fig. 13. The terminal cells in the pileipellis of the epitype were highly variable in form and dimensions, from 15 to >70 µm long, and not always cylindrical, but more irregular in form, particularly closer to the pileus margin, more or less similar to the pileipellis of *R. brunneoannulata*.

These Malagasy collections of *R. radicans* are unmistakable because of the bright yellow-orange color of the very thin, dull and farinaceous surface of the strongly striate-tuberculate pileus, the immediate appearance of bright orange-yellow stains after the slightest injury, the presence of an annulus and the “more-than-usual” number of shorter lamellulae and forked lamellae when compared to most other species groups in subgen. *Heterophyllidiae*.

**Russula radicans forma miomboensis fo. nov.** — Figs. 15-16

**Fig. 15.**
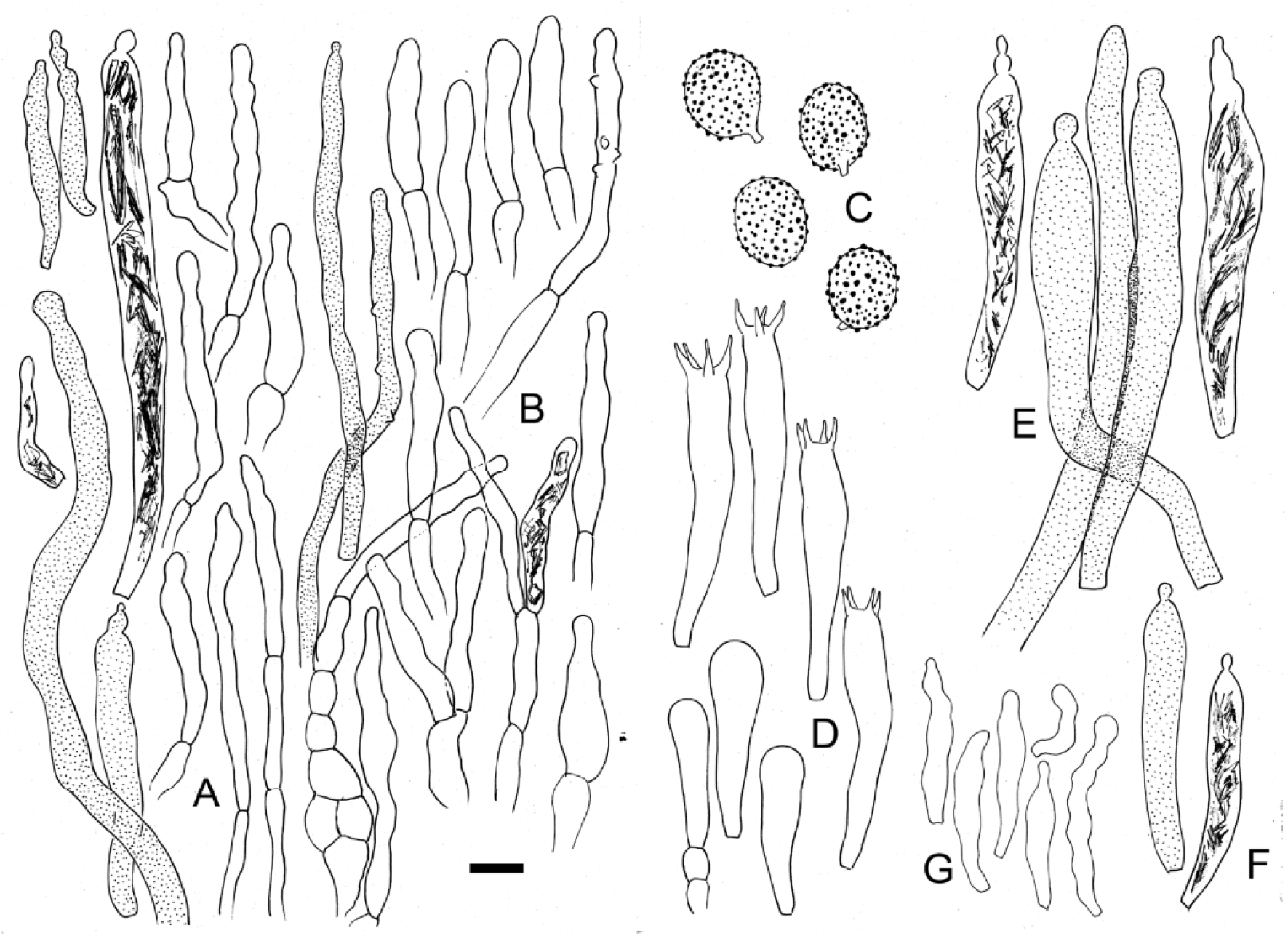
***Russula radicans* forma *miomboensis*** (holotype, BB4833) A. Long cystidioid hyphae from subpellis and shorter pileocystidia from near the pileus margin surface. B. Hyphal extremities and some pileocystidia from the pileus center. C. Basidiospore ornamentation. E. Basidia and basidiola. F. Pleurocystidia. G. Cheilocystidia. H. Marginal cells. Scale bar = 10 µm but only 5 µm for basidiospores. Drawings: B. Buyck. Drawings at 2000

**Fig. 16.**
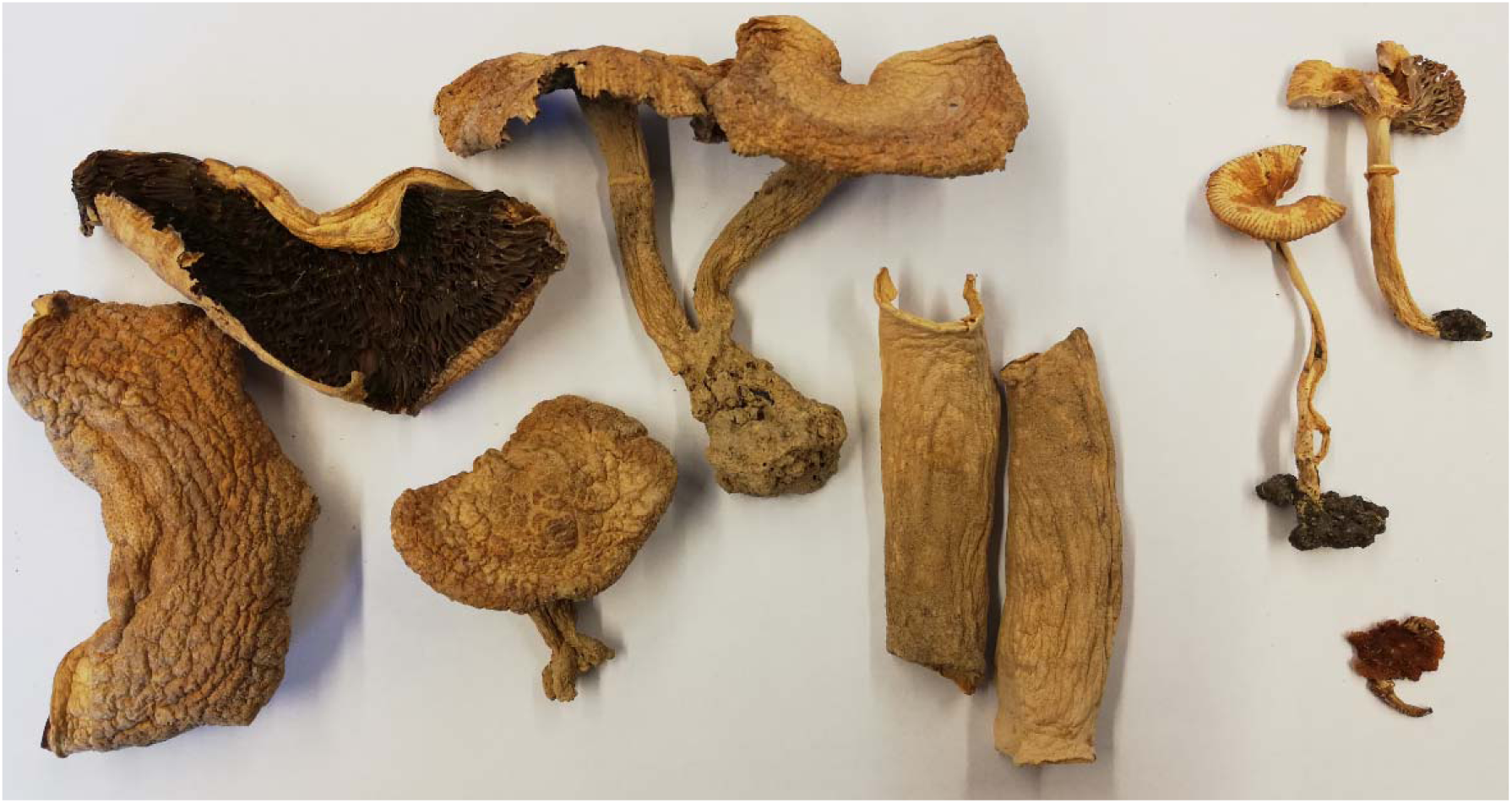
*Russula radicans*. Size comparison between dried basidiomata of the fleshy and large fo. *miomboensis* with dark chocolate brown lamellae (a, Buyck 96.019 from *Brachystegia* woodland in Zambia) on the left, and those of the Malagasy *R. radicans* fo. *radicans* epitype (upper right) and the *R. acriannulata* holotype (lower right). Scale bar = 1 cm

**Fig. 17.**
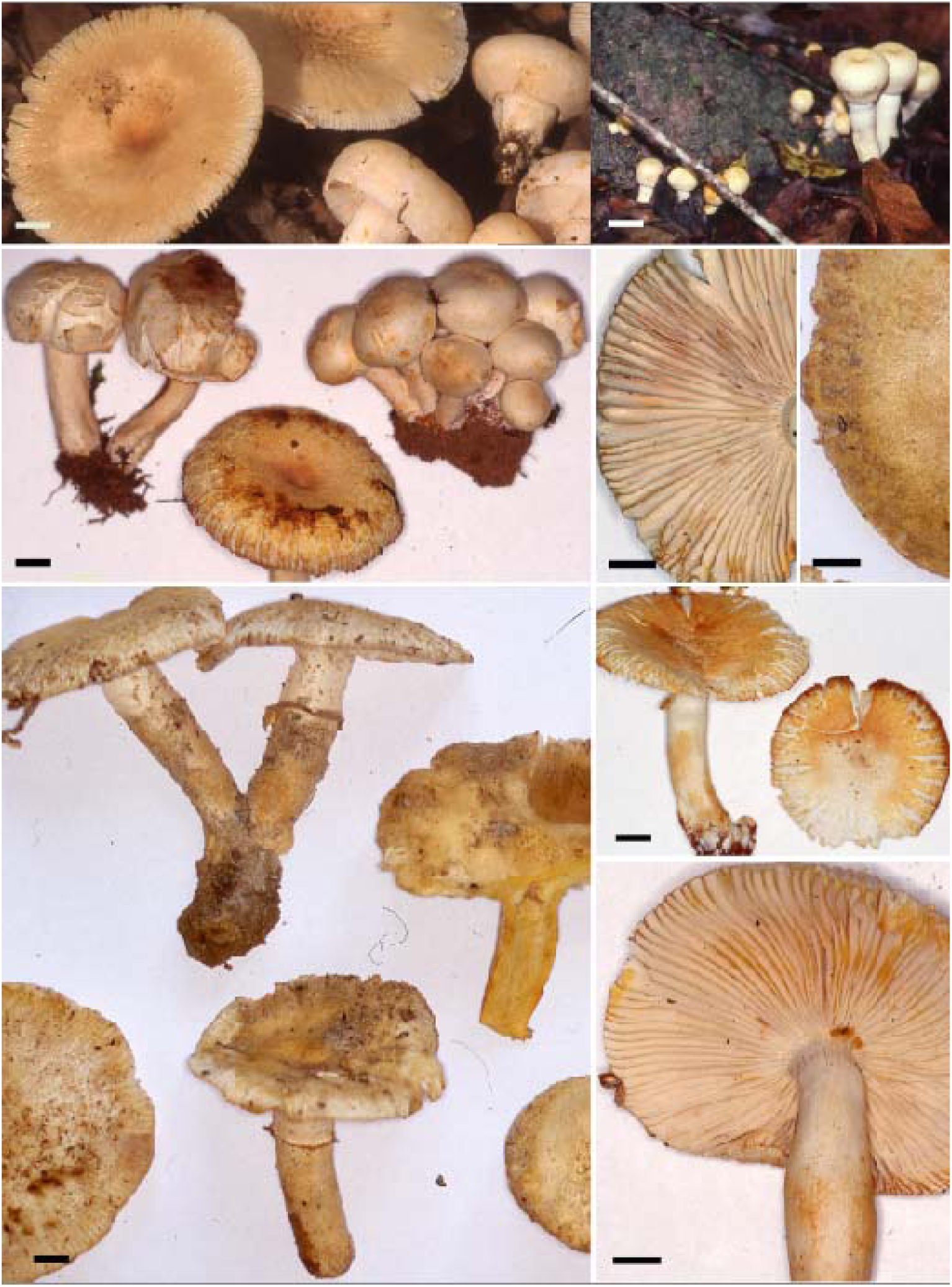
***R. radicans forma miomboensis*.** A. A collection from richer miombo woodlands. in Zimbabwe (D. Arora 99.1). B. Primordia fruiting on a dead trunk. C. Specimens collected from Brachystegia woodland in Burundi (BB4430). D. Detail of hymenophore (BB4430). D. Detail of pileus surface (BB96.019) D. Pale but strongly yellowing specimens in rich Zambezian woodland (BB97.019). E-F. Part of the holotype collection from dry, rocky *Brachystegia microphylla* woodland in Burundi. Scale bars = 1 cm. Photo credits: D. Arora (A), B. Buyck.

*Diagnosis*: although genetically identical to the typical form from Madagascar, it differs morphologically in the field in the much more fleshy and much larger fruiting bodies, the largest ones having lamellae discoloring dark chocolate when dried; microscopically, it also has less inflated cells in the pileipellis.

*Holotype*: **Burundi**. Bururi province. Nkayamba, in *Brachystegia microphylla*-woodland just N of Rumonge, alt.± 850 m, 18 December 1992, B. Buyck 4833 (PC0125146)

*Mycobank*: xxxxxxxx

*Etymology*: referring to the local name of the typical woodland habitat.

**Basidiomata** varying from large to medium sized, from fleshy and firm to very fragile, all parts turning rapidly to almost instantly bright yellow orange at handling but slowly so with age, terricole and usually forming large groups of up to > 100 fruiting bodies, sometimes fasciculated at the stipe base. **Pileus** (20)42–85(120) mm diam., regular, at first almost globose with an more or less flattened to depressed upper part, then becoming plane and gently depressed in the centre; pileus margin adnate to the stipe when young, then disrupting, leaving an annulus remaining attached to the stipe or pending from the pileus margin, becoming shortly striate-tuberculate with age; pellis entirely separable, dull, dry, during expansion of the pileus fissuring radially from the margin inwards and resembling sometimes veil-like remnants, entirely pale cream then becoming yellowish orange (5D6–7, 6A3) from the centre outward, yellow to cream towards the margin, finally orange yellow when old. **Stipe** central, 34–68 × 6–12 mm, sometimes annulate, subcylindrical to distinctly tapering towards the base, often distinctly sinuous-undulate or curved in the lower half, dry, white to cream (4A2) above the ring, becoming entirely yellow to brownish orange from the stipe base upwards, lacunar then spongy-fistulose. **Annulus** distinct in mature specimens, either remaining attached to the pileus margin or hanging freely around the stipe. **Lamellae** shortly adnate, up to 7 mm high, normally spaced (ca 1L/mm), white to whitish cream or even pinkish in young unopened caps, with occasional lamellulae or forking at various distances from the stipe attachment ; edge sometimes finely fimbriate when young, concolorous, rapidly staining yellow when touched. **Flesh** brittle, up to 6 mm thick in the pileus but often also very thin, in the pileus first whitish, then becoming yellowish orange (5A5), in the stipe first pale yellow then orange-yellow (4A6), especially near the base and in the cortex. **Taste** mild or nearly so. **Odour** weak, not disagreeable. **Spore print** whitish (0–1 Dagron Code).

**Exsiccatum** almost entirely pale greyish to yellowish cream (4A2, 4B2–3) in young specimens, older ones usually greyish brown (5D2–5) in the pileus center, lamellae in larger specimens turning dark chocolate brown. **Macrochemical reactions** on stipe context: FeS0_4_: greyish green (27E6); Ammonium: greyish brown (5F3); KOH: dark orange (5A4–8); Phenol: greyish green to brown (7E6).

**Basidiospores** shortly ellipsoid, 7.53–***8.12–8.66***–9.77(9.97) × 5.93–***6.41–6.97***–7.86 µm (Q=l.15–***1.22–1.28***–1.35; n=5×20); ornamentation of relatively spaced, isolated, obtuse warts, up to 1 µm high; suprahilar spot not amyloid. **Basidia** elongate, (47)57–63 × 8–11 µm, four-spored ; sterigmata stout, 3–5(7) × 1.5–2 µm. **Hymenial gloeocystidia** abundant, slightly emergent, very variable in dimensions, near the edge sometimes as small as 25 × 5 µm, elsewhere usually voluminous and in some cases very long, 70–150(250) × (5)8–15 µm, running sometimes over considerable distance in the lamellar trama parallel to the hymenium (and therefore perhaps more reminiscent of pseudocystidia), thin-walled, cylindrical to subclavate, at the tip moniliform, capitate or simply rounded, less frequently narrowing to appendiculate; contents very prominent, coarsely crystalline to amorphous, SV +, brown in KOH. **Marginal cells** not particularly differentiated, very small, usually 20–35 × 5–7 µm, basidiolomorphous or irregularly inflated and ± elongate, mingled with abundant cystidia, resulting in a practically sterile lamella edge. **Lamellar trama** almost entirely composed of sphaerocytes and invaded by the deeply rooted bases of hymenial cystidia. **Pileipellis** ± 250– 300 µm thick in the pileus center, not well delimited from the underlying trama and obscurely two-layered, orthochromatic in cresyl blue, throughout mixed with very numerous, long pileocystidia and cystidioid hyphae. Hyphal extremities forming a dense mat on the pileus surface, mostly composed of very short shains of 2–3 subcylindrical of irregularly inflated cells up to 7 diam., very large inflated cells in lower layers absent, but smaller more or less globose or ellipsoid cells – sometimes in chains - present and up to 10 µm diam.; the terminal cell (26)36–61(74) × 6–8(9) µm, usually slender and subcylindrical, sinuous in outline or repeatedly slightly constricted, tapering near the tip or ± capitate, rarely fusiform, lageniform or variously inflated, diverticulae frequent..Pileocystidia very abundant, those placed terminal on the hyphal extremities in the suprapellis mostly 25–40(60) × (3)5–6(10) µm, those emerging from deeper layers up to several 100 µm long and 6–10 µm wide; contents abundant, coarsely crystalline, SV +. **Clamp connections** absent in all parts.

*Additional examined material*: **Burundi**. **Bururi province**. Nkayamba, in *Brachystegia microphylla*-woodland just N of Rumonge, alt.± 850 m, Dec.14, 1991, Buyck **4017** (PC0125131); ibid., Dec.18, 1991, Buyck 4036 (PC0125132); ibid., Dec.26, 1991, Buyck **4152** (PC0125134), Buyck **4156** (PC0125133), Buyck **4158** (PC0125135); Buyc **4360** (PC0125140); ibid., April 29, 1992, Buyck **4430** (PC0125141), **4440** (PC0125142), **4447** (PC0125143); ibid., Dec.18, 1992, Buyck **4781** (PC0125144), **4790** (PC0125145),; ibid., 18 Dec. 1994, Buyck 5627;(PCxxxx) Nyamirambo, Rumonge Forest Reserve., in recolonizing *Uapaca*-*Brachystegia* woodland (after several years of manioc cultivation), May 18, 1993, Buyck 5131 (PC0125147), Buyck **5327** (PC0125150), Buyck **5334** (PC0125151), Buyck **5371** (PC0125152), Buyck **5376** (PC0125153). **Cankuzo prov**., under *Julbernardia globiflora*, 11 March 1994, leg. B. Nzigidahera, Buyck 01.008 (PC0125156). **Zambia**. **Copperbelt province**. Gibson’s farm, in degraded miombo woodland, leg. Buyck & Eyssartier, 16 Jan. 1996, Buyck 96.019 (PC0125157); near Chibuli, in miombo woodland, leg. Buyck & Eyssartier, 28 Jan 1996, Buyck 96.213 (PC0125158). **Zimbabwe**. Near Chisipite, close to Harare, 12 Jan 1999, D. Arora 99-1.

*Notes*: It was quite a surprise to obtain identical or near-identical ITS sequences (Fig. 2) for the collections from the Zambezian miombo woodlands of Burundi, Zambia and Zimbabwe, when comparing their overall habit with the much smaller, slender and differently coloured specimens from littoral rain forest in Madagascar (see Fig.14). Based on our experience in Africa, we would suspect that *Russula* species from the humid littoral forests on Madagascar’s east coast have their closest relatives in the Central African rain forest area, not in the seasonal surrounding miombo woodlands and certainly not producing identical ITS sequences.

Compared to the Malagasy epi- and holotype of *R. radicans*, these miombo collections were associated with different host trees growing in different soils under different climatic conditions, even when comparing the various miombo sites among them. The collections from the richer types of African miombo woodlands differ significantly in the field from the Malagasy holo- and epitype not only by their much larger size and much firmer, more fleshy context, but also by their broader lamellae (only± 3 mm in the Madagascar collections) that turn dark chocolate brown on drying, producing exactly the same colour as in fresh *Lactarius chromospermus* Pegler which also occurs in the Zambezian woodlands (see Buyck & Verbeken 1995). Nearly all collections from richer miombo areas are also often much paler, producing cream coloured to almost whitish fruiting bodies. Compared to the holo- and epitype from Madagascar that have an almost smooth, farinaceousand brightly coloured pileus surface, the areolate aspect of the pileus surface in fleshy specimens in the miombo collections is another conspicuous difference. Under the microscope, however, we were unable to find significant differences. The description of a simple ‘forma’ allows to draw attention to these very different overall aspects of the morphology within a single species.

Three years of weekly field work in Burundi demonstrated that *R. radicans* forma *miomboensis* appears shortly after the first rains in early November. Until the end of December, it forms rich fructifications, often in fairy rings around the base of trees and composed of a few to nearly a hundred basidiomata. Then, the species starts to disappear, but reappears again after the short dry season (somewhere in January-February), but in much smaller numbers. It occurs in the Zambezian miombo woodlands most likely associated with various Fabaceae. In Burundi, we found it to be particularly common in very exposed, hilly and rocky areas under *Brachystegia microphylla*, but the species is also frequent under the much larger trees of *B. bussei* or *B. utilis* in richer miombo types or in *Julbernardia globiflora* dominated miombo. In these miombo woodlands, this form of *R. radicans* can very easily be confused in the field with the equally robust *R. oleifera*, which can be very similar in size and overall colour and can also reproduce prolifically, forming groups composed of many fruiting bodies.

***Russula radicans*** forma ***sankarae*** (Sanon & Buyck) Buyck, ***comb. et stat. nov*.**

Basionym**: Russula sankarae** Sanon & Buyck, *Cryptogamie, mycology* 35(4): 389. 2014

*Diagnosis:* differs from *R. radicans fo. miomboensis* in its more fragile habit, the salmon pinkish colours on pileus, annulus and stipe and in being endemic to the gallery forests of West Africa.

*Holotype:* **Burkina Faso**. Province of Kénédougou, near the village of Dan, in gallery forest with *Berlinia grandiflora*, *Afzelia africana* and *Malacantha alnifolia*, N 10°54’01’’ – E 4°51’00’’, alt. 466 m, 11 July 2014, Sanon E. 192 (Holotype PC0142188, isotype OUA).

*Mycobank***: xxxxxxxxx**

**Pileus** (28)37–50(70) mm, thin-fleshed, brittle, margin shortly striate-tuberculate over 4– 17(25) mm depending on pileus size; cuticle not peeling, deep orange, brownish orange (7C4) to reddish orange (6–7B4–5) in the center, becoming salmon pink to pinkish orange (6–7A2– 4) towards the margin; dull, smooth and continuous in the center but dissociating rapidly into appressed squamules toward the margin during expansion, revealing the white colour of the flesh below, discolouring yellowish-brown to rusty brown with age or when injured.

**Lamellae** adnate to shortly decurrent, almost equal to unequal and intermingled with frequent lamellulae of variable length, bifurcations principally closer to the stipe, strongly turning orange-yellow when injured, not densely spaced (7–10L+l/ mm), 6–7 mm in height, white to pale cream; lamella edge smooth, concolourous. **Stipe** (25)35–47(66) × 7–15 mm, subcylindrical but usually tapering toward or near the base; surface white, but when young always with salmon pinkish tinges, especially in its lower halve, rather firm, becoming rapidly multi-chambered inside, then hollowing with age. **Annulus** very fragile, salmon pink, easily detaching from the stipe and hanging in fragments on the pileus margin. **Flesh** brittle, thin, ca. 1 mm thick at mid-radius, white strongly orange yellow (4A6–7) with age. **Odour** unremarkable. **Taste** mild. **Spore print** not obtained, probably white. **Macrochemical reactions**: FeSO4, salmon reaction in white parts of stipe; Sulfovanillin, KOH, Phenol, Guaiac all negative on stipe.

Microscopic features similar to forma *miomboensis* (for a detailed description, see Sanon et al. 2014)

*Examined material*: **BENIN**. Kossoucoingou, N 10° 09.859’ E 001° 12.084’, 507 m asl, on rocky soil under *Isoberlinia tomentosa*, leg. Yorou, 20 July 2021, CM 21.159/SYN5083 (B 70 0105449); Chutes de Kota, N 10°12’43.5” E 001°26’36.6”, 501 m asl, near riverbed in sandy soil under *Uapaca guineensis* and *Berlinia grandiflora*, leg. Manz, Hampe, Rühl, Sarawi & Dongnima, 15 June 2022, CM 22.177 (B 70 0105451); ibid., leg. Manz & Hampe, 05 July 2022, CM 22.278 (B 70 0105452); ibid., N 10° 12.769’ E 001° 26.758’, 503 m asl, leg. Yorou, 19 July 2021, CM-21.161/SYN5072 (B 70 0105450); ibid., N 10°12.659’ E 001°26.833’, on rocky soil under *Isoberlinia doka, Monotes kerstingii, U. guineensis*, 498 m asl, leg. Manz, Hampe, Yorou, Gildas, 14 July 2021, CM 21.126 (B 70 0105447); ibid., N 10° 12.769’ E 001° 26.758’, on rocky soil under *U. guineensis, B. grandiflora*, 503 m asl, leg. Manz, Hampe, Yorou, Gildas, Daouda, 19 July 2021, CM 21.140 (B 70 0105448).

*Notes*: This new form was originally described as a different species from Burkina Faso in West Africa (Sanon et al. 2014). The species was described as new because of its unique and distinct pinkish to salmon tinges, the strong fragmentation of the pileus cuticle and its different distribution and host associations (although still including family Fabaceae) compared to the typical form from Madagascar.

In this paper, we report on new collections from Benin, southern neighbour to Burkina Faso. All these West-African collections were gathered in gallery forest as was the case for the holotype. The latter habitat is some sort of extension of humid forest along rivers in the more seasonal woodland area. These new collections confirm the presence of distinct pinkish tinges at least when young. Usually the entire young pileus is salmon pink when still closed, but the pink colour generally remains present after expansion, particularly near the pileus margin, as well as on the annulus and even on lower parts of the stipe. However, the here newly described *R. cameroonensis* (a close relative of *R. brunneoannulata*, see Fig.2) from the Central African rain forest shares with these West African collections the distinct salmon pink tinges and fragile, thin-fleshed habit, but both species are genetically quite different (15 to 16 base pair changes in the ITS, or slightly over 97% similarity) and also differ in their spore ornamentation which is distinctly lower in all forms of *R. radicans*.

All these West African collections share a single base pair difference (at position 508, T instead of C) in their ITS sequence compared to the other African or Malagasy collections of *R. radicans*, notwithstanding the sometimes important differences in general habit. From the Malagasy epitype of *R. radicans* it additionally differs in merely 1 bp (G/A) out of the 1300 sequenced base pairs in the LSU sequence, and also in 2 bp in the coding part of *tef1*, but 6 bp in the introns (excluded from our analysis), while *rpb1* and *mtSSU* sequences were identical. We, therefore, reduce *R. sankarae* here to a simple colour form of *R. radicans* that is associated with different host trees and endemic to West Africa.

**Fig. 18.**
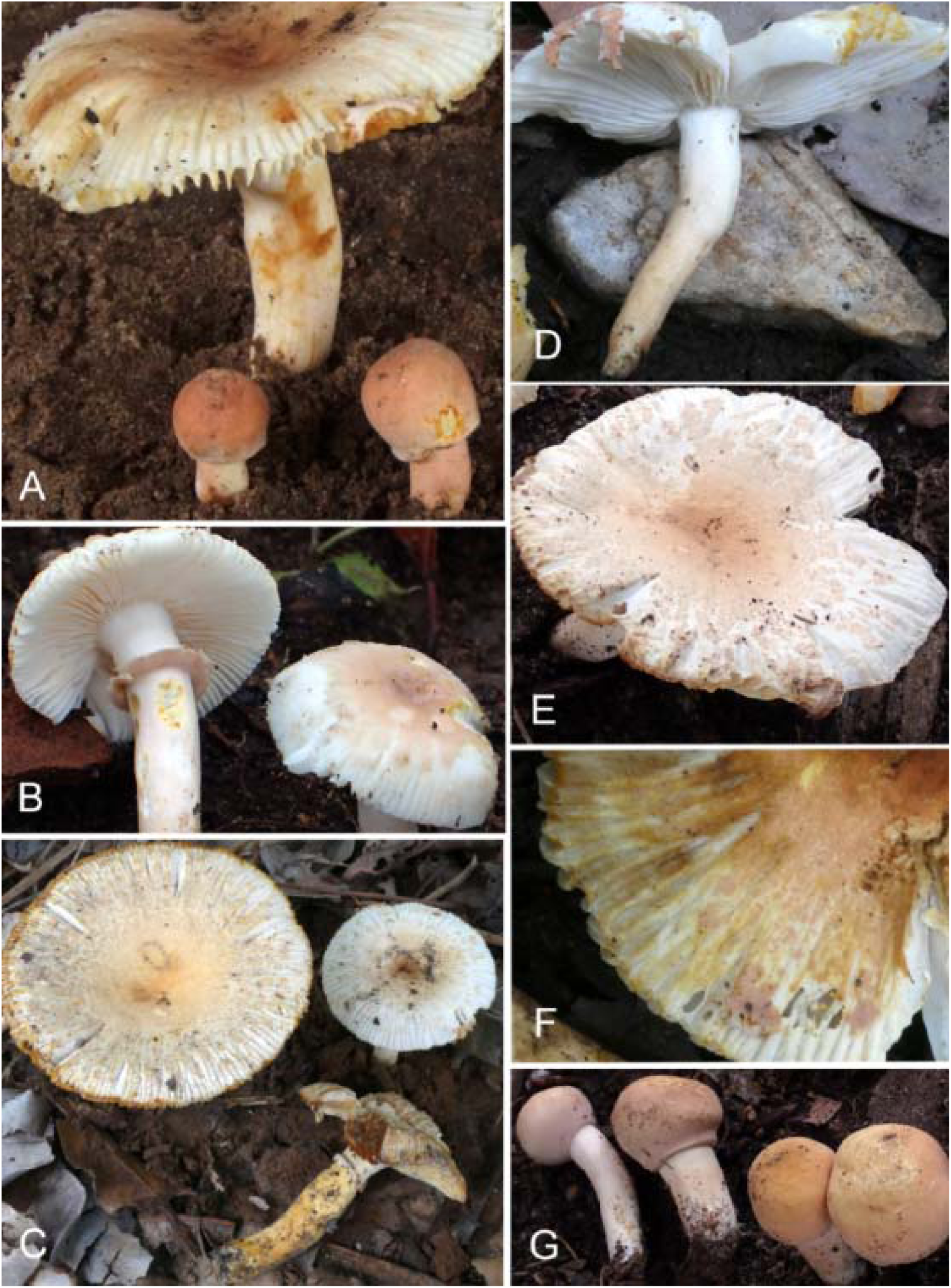
***Russula radicans*** forma ***sankarae***. A. Fruiting body with primoridia (CM 21.140). B. Fresh fruiting bodies in excellent condition; notice the unequal gills (CM 22.177). C. Unusually pale specimens (CM 21.159). D. Annulus (CM 21.126). E. Pileus with fissuring margin (CM 22.278). F. Detail of discolouring pileus surface (CM 21.126). G. Young primordia (CM 22.278). (photo credits: C. Manz & F. Hampe)

***Russula tapiae*** Buyck & Randrianjohany s**p. nov**. — Figs. **19**-**20**

**Fig. 19.**
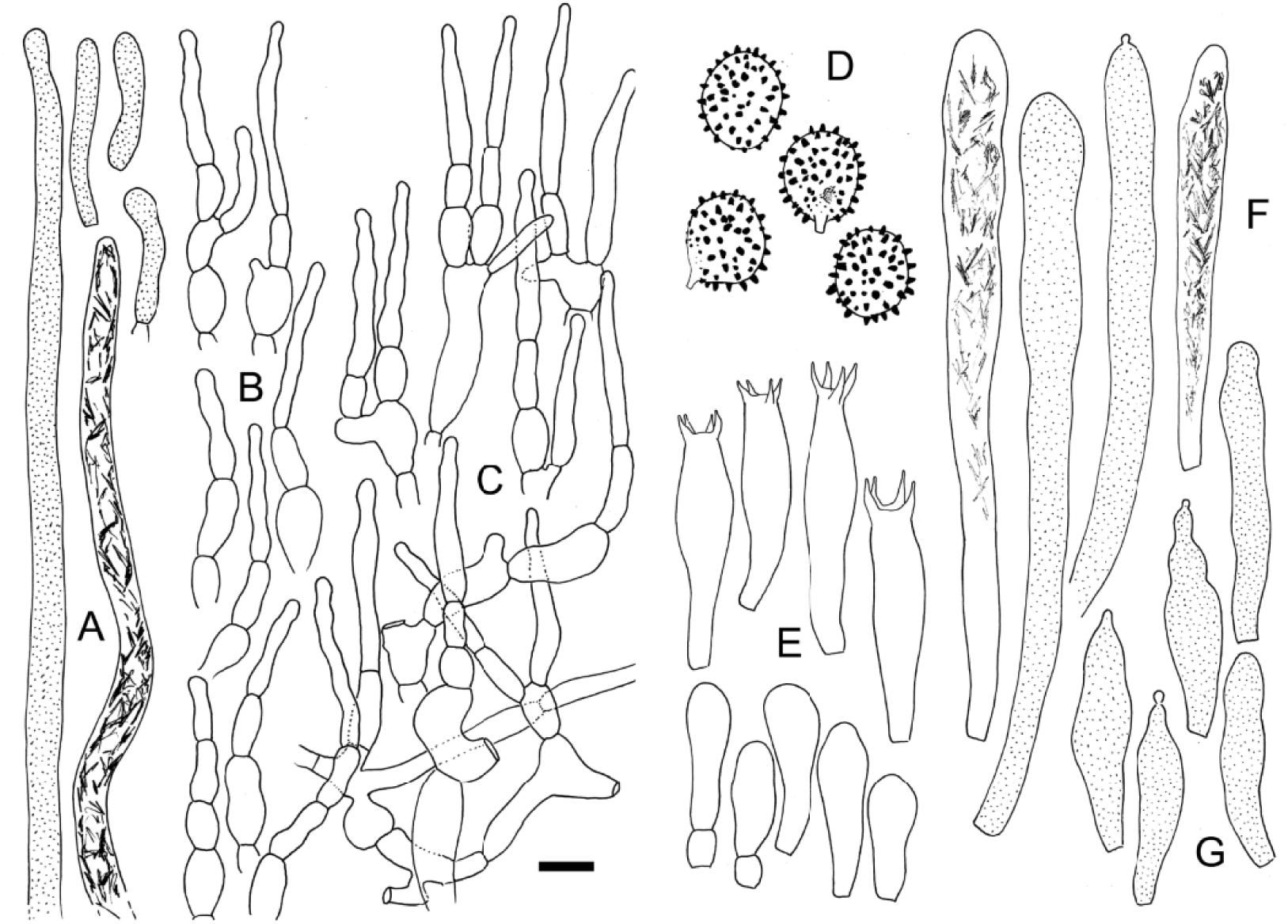
***Russula tapiae*** (holotype).A. Cystidioid hyphae from subpellis and shorter pileocystidia from the pileus surface. B. Hyphal extremities from the pileus center. C. Hyphal extremities from the pileus margin. D. Basidiospore ornamentation. E. Basidia and basidiola. F. Pleurocystidia. G. Cheilocystidia. Scale bar = 10 µm but only 5 µm for basidiospores. Drawings: B. Buyck.

**Fig. 20.**
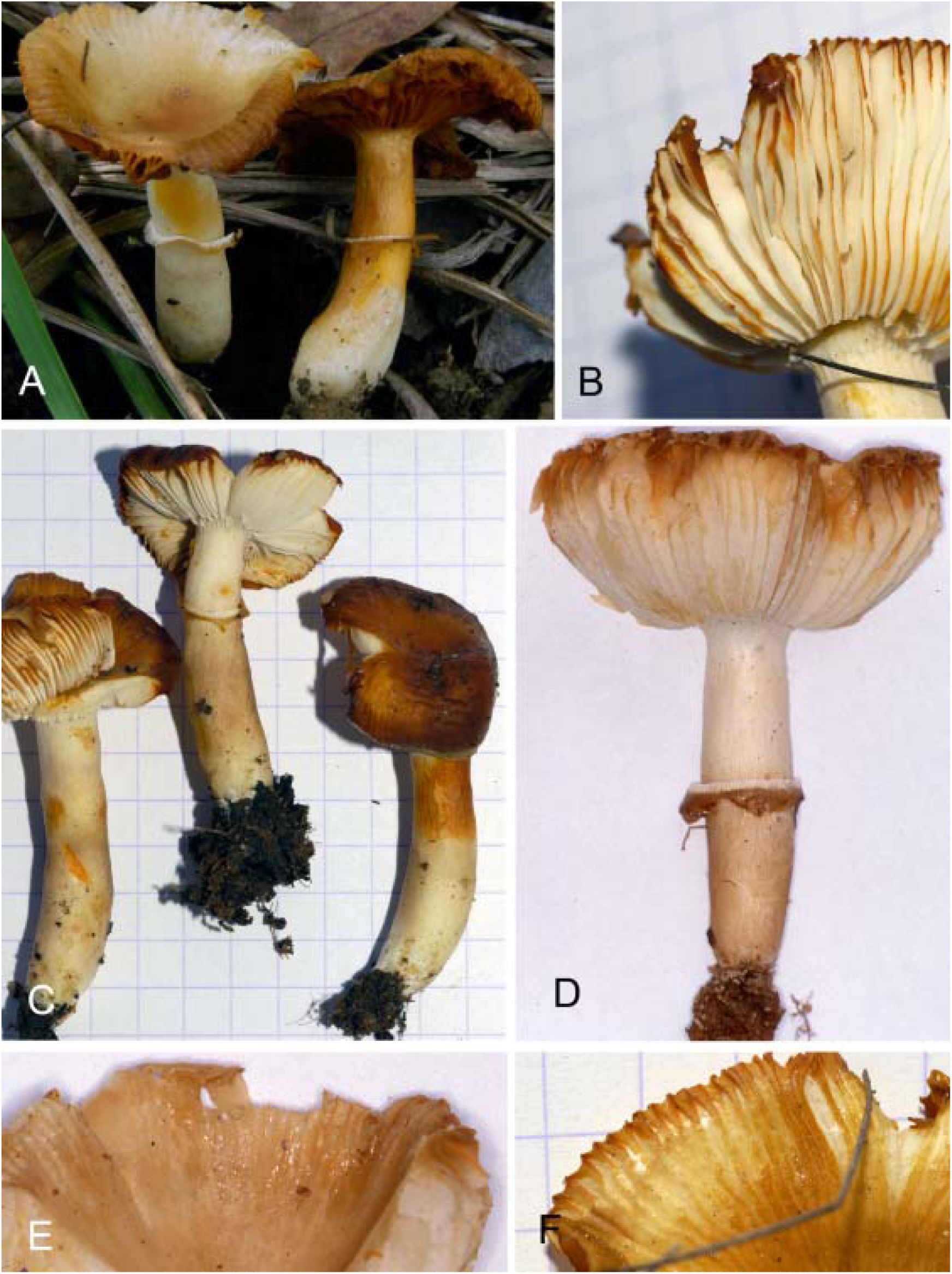
***Russula tapiae.*** A. Field habit (holotype). B. Detail showing lamellae mixed with lamelllulae (Buyck 08.322). C. General habit (Buyck 08.322). D. General habit (BB97.109). E. Detail of pileus surface (BB97.109). F. Detail of pileus margin (BB08.322). Photo credits B. Buyck.

*Diagnosis*: differs from other species in the subsection by the combination of the pale stipe, its association with the endemic *Uapaca bojeri* woodland in Central Madagascar and the rapid cinnamon brown discoloration of the tissues when old.

*Holotype*: Central Highlands, 50 km South of Antsirabe along N7, on the “Col des Tapias”, 1500 m asl., in degraded *Uapaca bojeri* woodland with intrusion of pine and eucalypts, 29 January 2008, Buyck 08.235 (PC0125171)

*Mycobank*: xxxxxx

*Etymology*: named after the local name ‘tapia’, which refers to the *Uapaca bojeri* woodlands

**Pileus** up to 38 mm diam., becoming strongly and widely depressed in the center, very thin-fleshed and fragile, surface completely peeling, dull to somewhat greasy, smooth and glabrous, continuous, mixture of pale brown, yellowish and pink, turning rapidly reddish brown with age, margin striate-tuberculate up to mid-radius. **Lamellae** adnate to subdecurrent with a tooth, subdistant (less than 1/mm), 6–7 mm high, unequal to subequal because of irregularly inserted lamellulae here and there, also forking sometimes, interstitial spaces with distinct transverse, concentrical anastomoses, slowly yellowing, then turning cinnamon brown; edges concolorous, often distinctly serrulate. **Stipe** central, 32 × 6–8 mm diam, cylindrical or often somewhat widening toward the base, white near the lamellae (above the annulus mostly) and at the extreme base, in between with ochre-orange to pinkish-rusty tinges, strongly yellowing where injured, particularly in basal part; stipe interior chambered with up to 6 cavities or more, finally hollowing. **Annulus** rapidly brownish at the distal end, after pileus expansion remaining either around the stipe or evanescent (detersile) with fragments temporarily attached to - or hanging from - the pileus margin. **Context** cream, very thin (ca. 2–3 mm above lamellae attachment) slowly yellowing, finally orange, than brown. **Odor** insignificant. **Taste** mild. **Spore print** not obtained, white on gill deposits.

**Macrochemical reactions**: FeSO_4_ on stipe: pale pinkish.

**Spores** ellipsoid and quite large in comparison with the other species of this clade, (7.50)7.86-**8.27**–8.68(8.96) × (6.46)6.75–**7.08**–7.42(7.71) µm, Q = (1.06)1.10–**1.17**–1.24(1.32), ornamentation not very dense, composed of strongly amyloid, isolated, cylindrical to conical spines, up to 1 (1.5) µm high, and smaller, obtuse warts; suprahilar spot verruculose or even weakly amyloid (greyish in Melzer) in its distal part. **Basidia** rather slender, 40–45(50) × 9.5–10.5 µm, clavulate or most inflated in their median part, 4-spored with sometimes stout sterigmata, 5-8 × 2 µm, near the lamella edge also 1–3-spored. **Basidiola** clavulate, slender. **Hymenial gloeocystidia** abundant, easily visible because of their yellowish contents, (in Melzer very yellow too) turning brownish in KOH, very variable in size, the longest ones originating from deep within the trama and up to 150 µm long, 11–12 µm wide, near the lamella edge with many smaller ones, mostly 27–37 × 5–8 µm, but also with longer ones that are rampant on the edge; all cystidia thin-walled, mostly clavate-pedicellate and with an obtuse-rounded or minutely mucronate apex, near the lamella edge also often fusiform; contents abundant, coarsely crystalline. **Marginal cells** not differentiated. **Subhymenium** pseudoparenchymatic. **Lamellar trama** mainly composed of large sphaerocytes. **Pileipellis** two-layered, orthochromatic in cresyl blue, subpellis ill-delimited from the underlying trama and composed of slender hyphae and abundant cystidioid hyphae and ascending pileocystidia; suprapellis in the pileus center forming a loose tissue of ascending and branching hyphal extremities that locally aggregate together in small tufts and arise from a thin pseudoparenchymatic layer; these extremities mostly consisting of 2 to 4 cells that are gradually getting narrower toward the terminal cell which is also distinctly longer, 19–36 × 2.5–4 µm, subcylindrical, sometimes slightly constricted subapically; the subterminal one shortly cylindrical to narrowly ellipsoid and often slightly constricted in its median part, lower cells more inflated, ellipsoid, 7–8 µm diam., and branching (usually with 1 or 2 other hyphal ends) from a much larger, often irregularly inflated cell; near the pileus margin essentially composed of similar hyphal extremities that are less organized in tufts and more spread out, without a pseudoparenchymatic layer and longer, more irregularly inflated basal cells up to >30 µm long and up to 15 µm wide. Pileocystidia cylindrical, obtuse rounded at the tip; those constituting terminal cells in the hyphal extremities often short, 25–35 × (4)5– 6 µm, in the subpellis much, much longer and up to 8 µm wide.**Cystidioid hyphae** in subpellis and underlying trama abundant and very apparent because of their yellowish, coarsely crystalline contents. **Oleiferous hyphae** not observed. **Clamp connections** absent.

*Additional examined material.* **Madagascar**. Central Highlands, 50 km South of Antsirabe along N7, on the “Col des Tapias”, 1500 m asl., in degraded *Uapaca bojeri* woodland with intrusion of pine and eucalypts, 3 February 2008, Buyck 08.321 (PC0125172), 08.322 (PC0125173), 08.323 (PC0125174); Arivonimamo, 1400 m asl, in *Uapaca bojeri* woodland, 30 January 1997, Buyck 97.109 (PC0125175). Vinany (Ambatofinandrahana), along the road to Itremu, under *Uapaca bojeri*, leg. Decary 20 Feb 1938, in R. Heim 6 (PC0714875); ibidem, under *Uapaca bojeri,* leg. Decary 19 Feb 1938, in R. Heim7 (PC0714876).

*Notes: Russula tapiae* is associated with the *Uapaca bojeri* woodlands on the Central Plateau in Madagascar and is widely distributed in this habitat but, considering the very few collections we made over the years, seems a rather rare species growing in groups of 2–3 individuals at the most. The two collections made by Decary in 1938 and sent to Paris were probably not studied by Heim and there is no mention of these in any of Heim’s publications. They were kept in alcohol which dried out over the years and are in a poor state.

Compared to *R. radicans* fo. *miomboensis*, (which is the only other taxon in the subsection that occurs in woodland habitats), *R. tapiae* is less energetically yellowing, but strongly and rapidly browning instead (the colour of coffee with milk or cinnamon). Microscopic differences concern mainly the higher spore ornamentation for *R. tapiae* and the hyphal extremities in the pileipellis that resemble much more the typical extremities found, for example, in species of subsection *Griseinae*.It shares these differences with its two closest relatives from the African rain forest, *R. cibaensis* and *R. afrovinacea*.

***Russula xylophila*** Beeli, Bull. Jard. Bot. Etat Brux. **14:** 84, Pl.2, fig. 1, -fig. 271, planche 70/3 (1936).— Figs. 21-22.

**Fig. 21.**
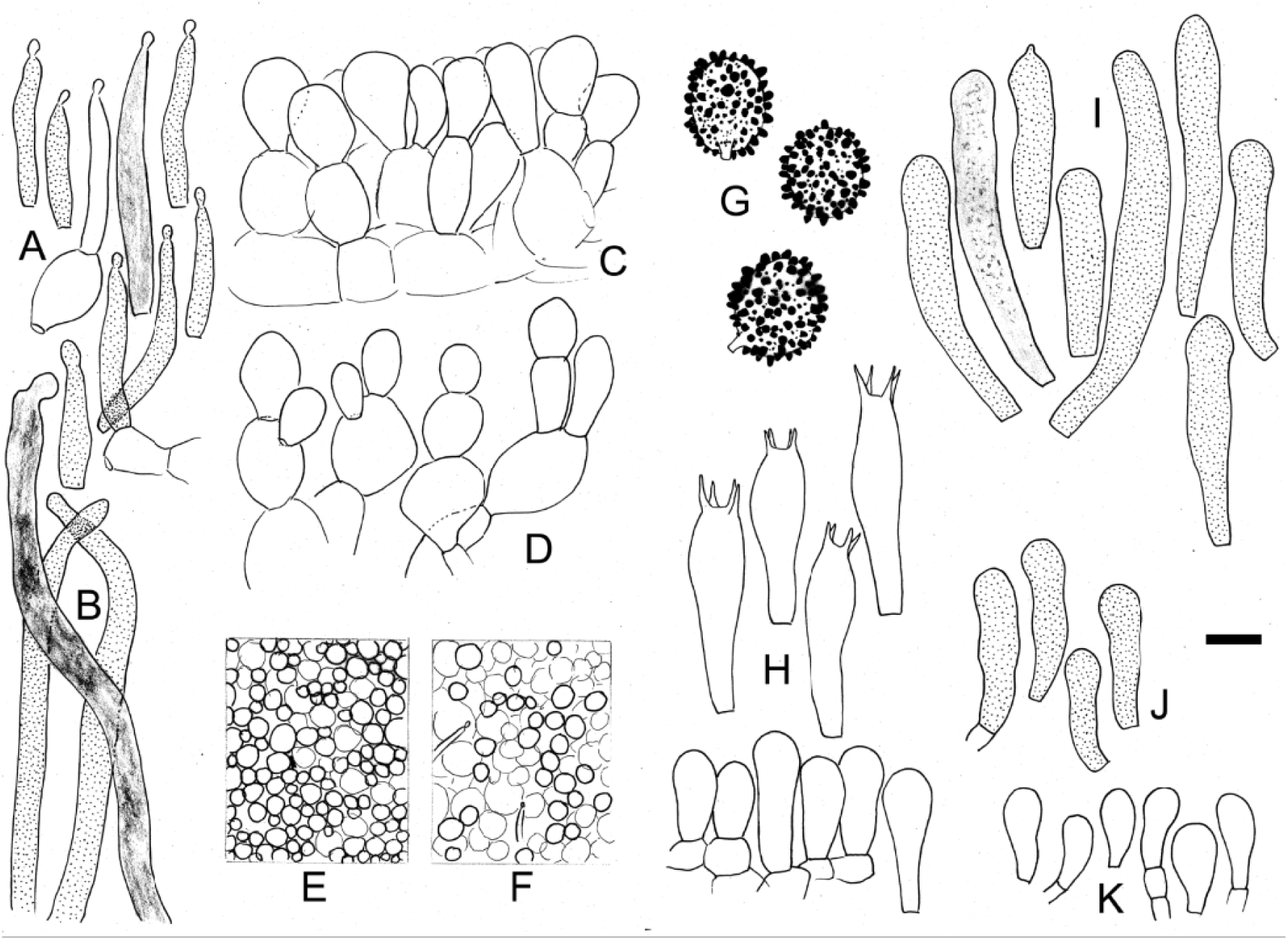
***Russula xylophila*** (epitype). A. Short pileocystidia at the pileus surface. B. Cystidioid hyphae from the subpellis and trama underneath. C. Fragment of a section of the suprapellis in pileus center. D. Hyphal extremities from the pileus margin. E. Surface view in pileus center showing most cells arriving at the same level. F. Surface view near pileus margin which is more uneven. G. Basidiospores observed in Melzer reagent. H. Basidia and basidiola. I. Pleurocystidia. J. Cheilocystidia. L. Marginal cells. Scale bar = 10 µm but only 5 µm for basidiospores. Drawings: B. Buyck.

**Fig. 22.**
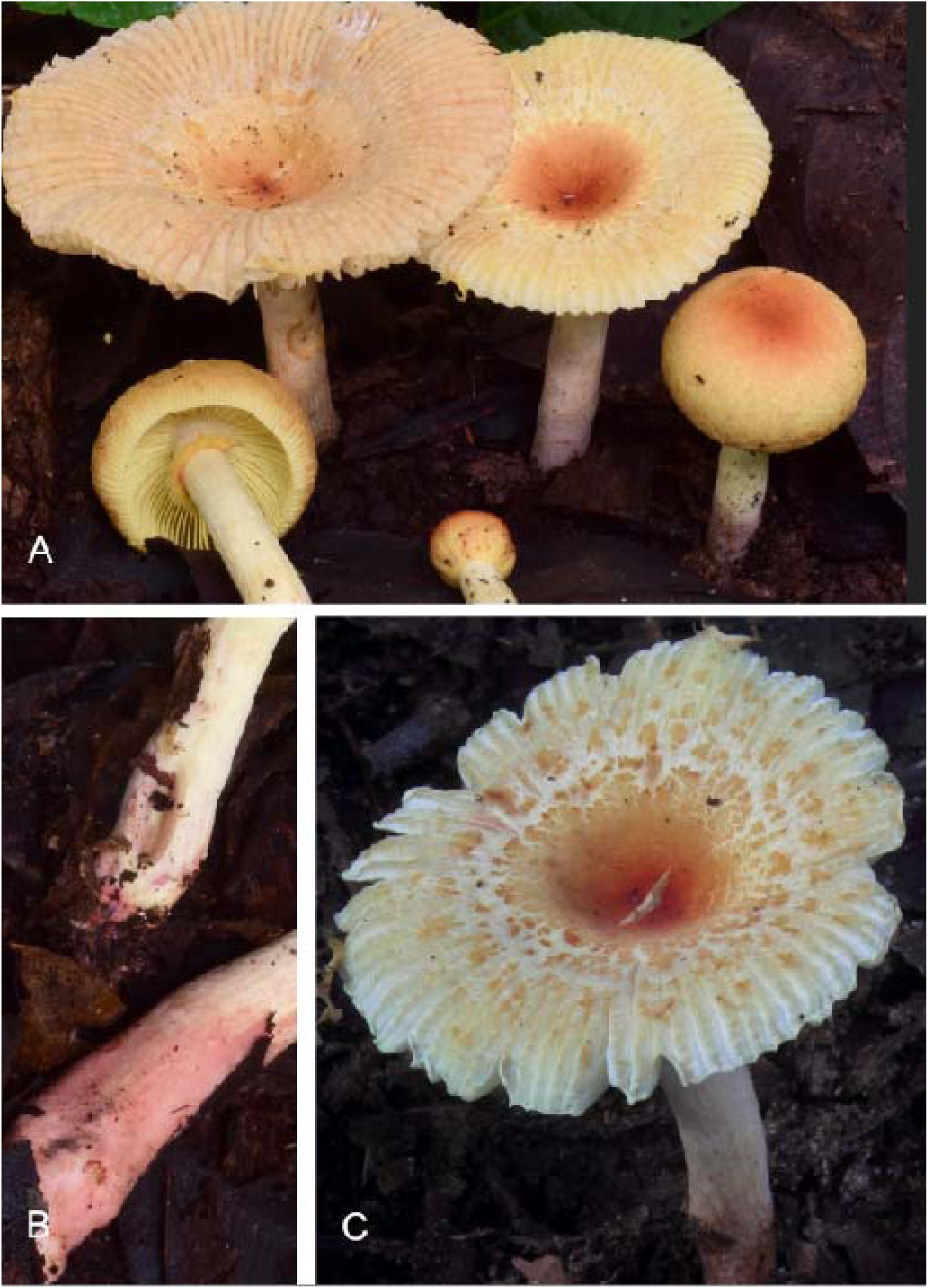
***Russula xylophila.*** Field aspect a. Notice the strongly striate-tuberculate pileus margin and distinct yellow tinge of the young lamellae (N. Siegel 2917). B. Reddish fibers on the lower stipe surface (T.W. Henkel 10433). C. Example of more contrasting colours of the fragmenting suprapellis (Dja96). Photo credits N. Siegel.

*Holotype*: **Democratic Republic of the Congo.** near Binga, on dead wood in seasonally flooded rain forest, December 1928, Goossens-Fontana 858 (holotype BR)..

*GenBank*: all new ITS sequences, also for the holotype

**Basidiomata** in groups, solitary, sprouting from between the dead leaves and other detritus on the forest floor. **Pileus** 18–63(83) mm diam., initially closed around stipe apex, flattened-convex with incurved margin in contact with the stipe, with expansion becoming broadly convex to planate with broad, shallow, central depression, covered with an orange red (7– 8B8) epithelial layer that progressively over the outer 2/3 of the pileus surface in a few larger fragments sitting on a yellow background, becoming deeply sulcate-striate and tuberculate, not peeling. **Lamellae** thin, adnate, up to 3–5 mm heigh, narrowing to 1.5 mm near extremities, subclose, subequal from frequent lamellulae, 2–30 mm long, irregularly inserted among lamellae (at least one every 2–3 lamellae), with some forkings present particularly closer to the stipe, pale yellow (2A3, 3A2–3)), unchanging from injury, but somewhat browning with age; edges concolorous, smooth to cystidiate and reddish brown. **Stipe** subcylindrical, either slightly narrowing or widening toward base, 35–70 × 6–8(17) mm, very pale yellow (2A2) over upper halve and densely tomentose, tinged more brownish over basal 1/3 to 2/3 and with patches of longitudinally oriented burgundy red (8D5) innate, repent fibrils that continue as basal tomentum at the extreme base; inside white, stuffed to chambered. **Annulus** mostly retained around the stipe, but also pending in separate fragments from the pileus margin, concolorous with pileus surface on the outer side, yellowish on inner side. **Context** pale yellow (as the young lamellae), later fading to pale cream, very thin (< 0.5 mm near pileus margin, 2 mm above stipe). **Taste** mild, perhaps slightly disagreeable or somewhat spermatic, but definitedly not acrid. **Smell** unremarquable. **Spore print** white in heavy deposit.

**Basidiospores** ellipsoid, (7.3)7.4–**7.82**–**8.24**–8.45(8.54) × (5.83)5.99–**6.33**–**6.48**–6.82(7.08) µm, Q = (1.16)1.19–**1.24**–**1.28**–1.35(1.38); densely to very densely ornamented with isolated or locally confluent, conical to short cylindrical warts, highly variable in size, some ponctiform and barely elevated, others up to 1(1.5) µm high and strongly amyloid, lacking ridges or linear elements; suprahilar spot verruculose, non-amyloid. **Basidia** 32–43 × 9–11 µm, tetrasporic, clavate; sterigmata medium-sized, 5–7 × 1–2 µm. **Hymenial cystidia** numerous,on lamellae sides (ca. 2000/m2), not very emergent and mostly rather short, generally 35–80 × 7–9 µm; near the gill edge sometimes as small as 22 × 7 µm, some rare cystidia much longer and arising from deep within the lamellar trama, generally subfusiform to clavate, mostly obtuse-rounded at the tip and without any appendages, thin-walled; contents very abundant, dense, granular-oily, distinctly yellow in KOH, blackening in sulfovanillin. **Marginal cells** unremarkable, very similar to young basidiola, but slender and more distinctly clavate. **Pileipellis** entirely orthochromatic in cresyl blue; subpellis composed of a weakly gelatinous layer of hyphae measuring approx. 1–3 µm in diameter, intermixed with very abundant cystidioid hyphae with yellowish, dense oily-granular contents, strongly blackening in sulfovanillin, mostly 3–7 µm wide, sometimes several hundreds of µm long, with clearly zebroid-encrusted walls, cylindrical, obtuse-rounded or somewhat narrowing at the apex; suprapellis a compact, dense layer of branched, short chains of not more than 3 to 5 swollen cells that become gradually smaller towards the surface, terminal cells usually 5– 12(15) µm diam., giving the impression of an uneven epithelium when seen from above, the lower cells voluminous, up to 25 µm diam., ellipsoid, globose or variously shaped, forming a ca. 50 µm thick pseudoparenchyma; pileocystidia 23–55 × 3.5–5 µm, rather few and dispersed, very inconspicuous (particularly in the pileus center) and emerging from among the voluminous articles of the pseudoparenchyma, slender, subulate to lageniform, minutely capitate, with similar yellowish, dense-oily contents. **Stipitipellis** producing very long trichoids of parallel hyphae having walls encrusted with amorphous deposits that are adhering to the lower stipe surface. **Clamp connections** absent.

Examined material: **Cameroon**. Dja Biosphere Reserve, within 300 m of base camp located at 3°21’29.8” N 12°43’46.9” W, in riparian forest under *Uapaca* sp., 19 August 2017, N.Siegel 2197 (PC0142591, HSC-F 004452); ibid., 21 August 2017, T.W. Henkel 10433 (PC0142590, HSC-F 004450); ibid., 7 September 2014, Dja96 (PC0142592, HSC-F 004451).

*Notes:* The original description (Beeli 1936) of *R. xylophila* was based on a single collection from the Central African rain forest. Later, Buyck (1989) reported two more microscopically identical collections, unfortunately lacking any notes or illustrations, from dry dense forest (locally referred to as ‘muhulu’) in Haut-Katanga, a province that lies in the wetter miombo zone, and is rich in riverine dense forests and dry evergreen forest patches (Meerts 2016). The presently studied specimens are the first documented collections of this species since its original description. Their identity was supported by an ITS sequence (GenBank PQ237116) obtained from the holotype collected in December 1928. However, since there is a clearly confirmed five base pair difference with the Cameroon collections as well as some microscopic differences (see below), we refrained from epitypification at this time.

These new collections confirm the overall picture sketched in the watercolours by Goossens-Fontana (in Buyck 1994). We can now confirm the white spore print mentioned in Goossens-Fontana’s original notes, but not the acrid taste (a recurrent problem in Goossens-Fontana’s descriptions due to many years of anti-malaria treatment), which was definitely mild in all new collections. Other adjustments to Beeli’s original description concern the presence of lamellulae and the clearly pale yellowish to even distinctly yellow colour of the lamellae when young (this was correctly shown in Goossens-Fontana’s watercolours of the species, but not mentioned in the description (see Buyck 1994). The basidiospores of the Cameroon collections are somewhat more ellipsoid and notably smaller than those of the holotype [HT: 8.4–**8.98**–9.5 × 7.1–**7.70**–8.3 µm (Q = 1.10–**1.17**–1.22)], and to some extent this also applies to the dimensions of basidia and hymenial cystidia.

*Russula xylophila* was placed in subsection *Aureotactinae* by Buyck (1989) based on the shared overall habit, the spore ornamentation composed of isolated warts, the presence of an annulus and the shared lignicolous fruiting habit with *R. radicans* from Madagascar. Notwithstanding these obvious similarities with the *Radicantinae* described above, our multilocus phylogeny (Fig. 1) clearly places *R. xylophila* now in Sect. *Ingratae*, thereby confirming Buyck’s earlier hesitation about its placement in *Radicantinae*. This hesitation was partly based on the absence of a distinctly yellowing context in *R. xylophila,* but especially on its well-developed pseudoparenchymatic suprapellis and on the flask-like pileocystidia. The latter are typical for some other species groups in sections *Heterophyllae* or *Ingratae*, but not for *Radicantinae* where pileocystidia are more cylindrical. The same type of flask-like pileocystidia are present in *R. oleifera,* but the latter lacks the distinctly pseudoparenchymatic suprapellis. At that time [before 1989], a pseudoparenchymatic suprapellis was never reported for species placed in *Ingratae*. However, in recent years, several Asian *Ingratae* possessing a pseudoparenchymatic pileipellis have been described in the *R. senecis* clade (Song et al. 2018, Gosh et al 2021). The placement in *Ingratae* of *R. xylophila* now also explains the presence of the innate, wine red fibrils on the lower stipe surface and stipe base of *R. xylophila*, another feature shared to some degree with several other *Ingratae* (see for example Song et al. 2018). The collection ‘Dja96’ (Fig. 21C) from Cameroon illustrates the sometimes high similarity with *R. oleifera* because of the coarsely squamulose pileus surface, although the latter species is quite more robust.

Except for the intense yellow discolouration of all tissues upon the slightest injury, *R. radicans* from the type locality in Madagascar resembles *R. xylophila* in the field. Both share the intense yellow-orange-red tints in the pileus center, the slender habit and the annulate basidiomata. *Russula xylophila* even has the yellowish tints in its lamellae when young. It is also the first annulate species that is firmly placed in sect. *Ingratae*. The only other annulate species which has been attributed to *Ingratae*, *R. cingulata* Buyck & E. Horak (1999) from Papua New Guinea, still needs molecular confirmation of its systematic placement.

There are also similarities with the much smaller and paler *R. acriannulata*, particularly the spore ornamentation seems near-identical when observed under the optical microscope, although the higher rods or warts are more obtuse in *R. xylophila* and are mixed with more very small interstitial warts. Both species differ considerably in the abundance and contents of pileocystidia and the composition of the pileipellis, as well as in the form of their hymenial cystidia

## CONCLUSION

The exact affinities of *R. aureotacta* remain uncertain at this time as the senior author and collaborators were unable to collect any *Russula* specimens that matched the original description, notwithstanding many years of fungal exploration in Madagascar. As a result, the correct interpretation of this species remains impossible because Heim’s original collections are in such bad condition that detailed microscopic observation is no longer possible. We were nevertheless able to measure the spores of the type collection showing these to be distinctly smaller than in any species of *Radicantinae* [*R. aureotacta* I39=H20; (6.04)6.39–**6.65**–6,90 × (5.00)5.19–**5.43**–5.66(5.83) µm, Q = (1.07)1.16–**1.23**–1.29(1.38)]. Heim also stated that, although the spore ornamentation of *R. aureotacta* is of a very similar type as in *R. radicans,* it has rare connections in between the warts, which sets it apart from all species described above. To this, we have to add that Heim had apparently no problem to distinguish between *R. radicans* and *R. aureotacta*, notwithstanding their many similarities. Because Heim cites other collections for *R. aureotacta* (all lost), we can discard the hypothesis that it was just an odd collection of *R. radicans* in which all specimens had already lost the annulus. Also the greenish shades mentioned for the pileus of *R. aureotacta* as well as the overall habit of its basidiomata sketched by Heim (1937) is different from *R. radicans* and allies. We, therefore, suggest that it is more likely that *R. aureotacta* does not belong to this species complex and that *Aureotactinae* should again be restricted to its type species, while *R. radicans* and close allies should be placed in a subsection of their own. Considering the inamyloid suprahilar spot mentioned for the spores of *R. aureotacta*, subgenus *Heterophyllidiae* remains the best hypothetical place for it.

**Fig. 23.**
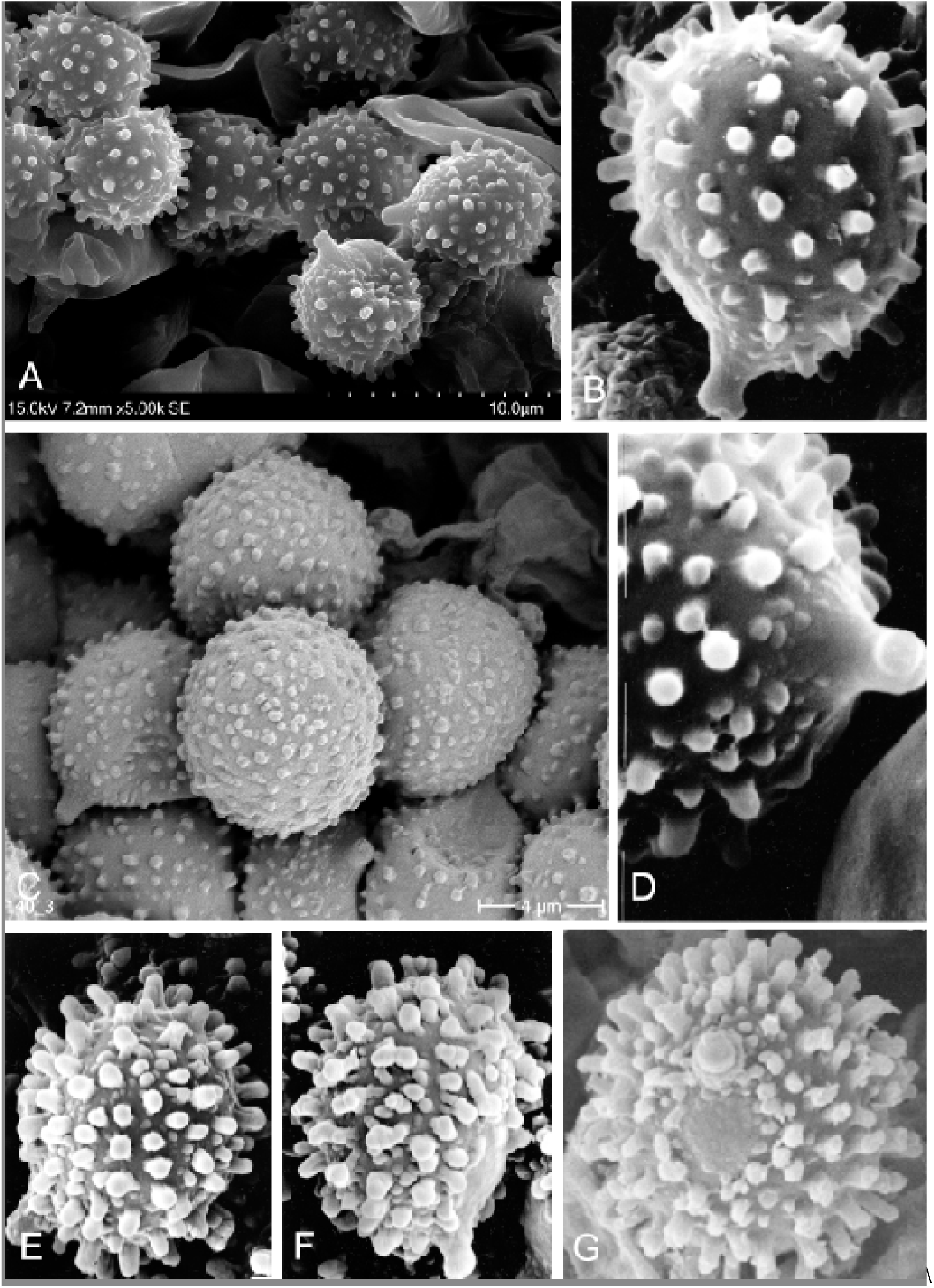
Different types of spore ornamentation among the discussed species. A-D. *Radicantinae*. E-G. Excluded *Radicantinae*.

**Subsection *Aureotactinae*** R. Heim ex Buyck, Bull. Jard. Bot. Nat. Belg. 60 :207(1990), emendavit Buyck

= *Aureotactinae* Heim, Prodrome a une flore mycologique de Madagascar et Dépendances, 1: Les Lactario-russulees du domaine oriental de Madagascar: 121 (1938), nom. inval.

*Diagnosis: Basidiomata* not annulate, centrally stipitate, pileus pale-coloured in tints of cream, pale yellowish or greenish, possessing abundant cystidioid hyphae and gloeocystidia throughout all tissues and producing spores with predominantly isolated warts and inamyloid suprahilar spot. Lamellae equal, white. Context yellowing, taste mild.

Type species: *R. aureotacta* Heim*, Rev. Mycol*. 2: 108 (1937).

**Subsection *Radicantinae* Buyck & X.H. Wang subsect. nov.**

= Radicantes R. Heim, Prodrome a une flore mycologique de Madagascar et Dépendances, 1: Les Lactario-russulées du domaine oriental de Madagascar: 125 (1938), nom. inval.

*Diagnosis:* **Basidiomata** lacking greenish or bluish tinges, usually very thin-fleshed and with strongly tuberculate-striate margin and a cuticle disrupting in smaller squamae outside the pileus center. **Lamellae** intermixed with more-than-usual lamellulae and occasional forkings at various distances from the stipe. **Annulus** present, often evanescent. **Stipe** multi-chambered inside, eventually becoming hollow. **Taste** mild or almost so. **Spore ornamentation** composed of isolated warts or spines, with verruculose, inamyloid or distally amyloid suprahilar spot. **Spore print** whitish. **Gloeocystidia** abundant in all tissues and filled with yellowish crystalline-oily contents provoking an intense yellow-orange discoloration upon injury as well as sometimes cinnamon brown discolouration of tissues or even a dark chocolate brown discoloration of the lamellae upon dessication.

*Type species*: ***Russula radicans*** R. Heim, *Candollea* 7: 389 (1938)

Mycobank: xxxxxxxx

Today, almost one century after the description of *R. radicans*, subsect. *Radicantinae* remains exclusively known from mainland Africa and Madagascar. As a result of this revision, the new subsect. *Radicantinae* holds six species and two more infraspecific taxa, as well as three more undescribed species that are only known from few environmental sequences (see Fig. 2).

Most *Radicantinae* are rare, with *R. cibaensis* from Madagascar and *R. afrovinacea* from Africa known from a single collection only. *Russula radicans*, originally described from the littoral rain forest in Madagascar where it is known from a single location, is by far the most common and most widely distributed species. Its range covers also the entire tropical African belt, from the Zambezian woodlands in East and Central Africa (forma *miomboensis*) to the Sudanian gallery forests (forma *sankarae*) in West Africa. Contrary to the rare species mentioned above, it is a frequent and profuse reproducer in the African miombo woodlands occurring often with dozens of fruiting bodies at a time. Additionally, being the only species in the subsection that occurs both in the seasonal woodlands and humid dense forest areas, *R. radicans* illustrates the enormous adaptability of a fungal species to the specificities of its habitat. It forms fruiting bodies that are extremely thin-fleshed, much more slender and up to a hundred times smaller in volume compared to those produced in the seasonal woodlands (Fig. 16A). *Russula radicans* is also a close relative of *R. brunneoannulata* and *R. cameroonensis*. All three species share a similar and original pileipellis structure that differentiates them from the other, much rarer species in the subsection which possess hyphal extremities much more similar to those typical for species in other subsections (e.g. *Griseinae*) of the subgenus.

*Radicantinae* represents an isolated lineage within subgenus *Heterophyllidiae*, but exhibits clearly a recent rapid diversification. The similarity among ITS sequences for all members of *Radicantinae* is 96% or higher, but then drops to < 87% for ITS sequences of the closest relatives (the asian *R. verrucospora* and american *R. redolens*). *Radicantinae* are morphologically most easily confused with several species in sect. *Ingratae* because of the often brownish tinges, the multi-chambered stipe, the strongly striate-tuberculate pileus margin, the abundance of gloeocystidia in all tissues and similar spore ornamentation. So far, the yellow-orange discoloration of their tissues appears to be the principal feature that differentiates them (but not from *R. aureotacta* whose systematic placement is still unknown). All *Radicantinae* share a similar configuration of the hymenophore, which is characterized by a “more-than-usual’ number of inserted lamellulae, too many to consider the lamellae as “equal”. A similar configuration can be found in some species of subsect. *Cyanoxanthinae* and in the few species that compose subgen. *Crassotunicatae* Buyck & V. Hofst., the sister clade of subg. *Heterophyllidiae* (Buyck et al. 2018, 2024). This is most probably a conserved feature, something in between the equal lamellae typical of the most recent clades in subgenera *Heterophyllidiae* and *Russula*, and the unequal or regularly forking lamellae characterizing the remaining, supposedly more ancient subgenera, as well as all of the other agaricoid genera in Russulaceae.

Host association is another interesting aspect of *Radicantinae* and clearly merits further investigation. The genus *Uapaca* (Phyllantaceae) is a remarkably frequent host or possible host tree for most of its species. In the Central African rain forests, *R. atrovinacea* was found under *Uapaca* sp., while *U. guineensis* is a confirmed host for R. sp1 from Gabon. *Russula radicans*, with its very wide distribution, has no molecularly confirmed hosts, but was collected in humid forests largely dominated by Fabaceae (*Intsia* in Madagascar and *Berlinia*, *Isoberlina* in West Africa), but *Uapaca* species were never far away (*U. guineensis* in W-Africa and *Uapaca* sp. in Madagascar). In the drier Zambezian region, we collected this species in woodlands dominated by various *Brachystegia* species, but also here *U. nitida* was a frequent intruder or even co-dominant tree. In Madagascar, *R. tapiae* is associated with *U. bojeri* dominated woodlands on the Central Plateau, while R. sp2 was collected in *Uapaca* dominated humid forests on the lower East coast. In the drier parts of the *Gilbertiodendron dewevrei* rain forest, the latter tree species is a confirmed ectomycorrhizal host for *R. cameroonensis* and also more than likely the single host for *R. brunneoannulata* based on field observations.

### Key to the accepted species

1a. Hyphal extremities of the pileipellis multicelled and slender, composed usually of more or less inflated and often branching basal cells giving rise to chains of 4 to 6 narrower cells, with the terminal cell usually quite longer but not wider than the subterminal cells (in other words hyphal extremities similar to species in e.g. subsect. *Virescentinae* or *Griseinae*) (2)

1b. Hyphal extremities of the pileipellis composed of mostly 2-3 cells of very variable form, with the majority of the terminal cells usually (much) larger and more inflated than the subterminal cells. (4)

2.a Pileus young reddish brown to vinaceous, rapidly much paler toward the margin during expansion, having a ring that remains attached in fragments to the pileus margin during pileus expansion. Under *Uapaca* in the Central African rain forest. Very rare.

***Russula afrovinacea***

2b. Pileus without reddish or vinaceous tinges, stipe annulate, endemic to Madagascar (3)

3a. Pileus pale yellowish brown, not strongly squamulose with age, turning cinnamon brown with age in all parts; occurring in the seasonal *Uapaca bojeri* woodlands on the Central Plateau. Widely distributed but rare.

***Russula tapiae***

3b. Pileus darker brown, becoming strongly squamulose-areolate toward margin; occurring in the humid forests of the Eastern escarpment. Very rare.

***Russula cibaensis***

4a. Spore ornamentation composed of low (0.5 µm), densely disposed, obtuse warts. (5)

4b. Spore ornamentation distinctly higher, composed of cylindrical warts or conical spines, but intermixed with some smaller, low warts, not very densely disposed. Occurring under *Gilbertiodendron dewevrei* in the Central African rain forest. (7)

5a. Occurring on the African mainland, in woodland or gallery forest. Common. (6) 5b. Occurring in littoral rain forest of Madagasxar’s East coast. Very rare.

***Russula radicans forma radicans***

6a. Basidiomata with distinct salmon pink tinges. Occurring in the gallery forests of West Africa.

***Russula radicans forma sankarae***

6b. Basidiomata without distinct salmon pink tinges, but variable in colour (cream, pale yellow, …). Occurring under Fabaceae in the Zambezian miombo woodlands.

***Russula radicans forma miomboensis***

7a. Basidiomata without pinkish tinges, stipe and pileus dark reddish brown over most of their surface; terminal cells of the pileipellis variable in shape, not principally fusiform. Locally common.

***Russula brunneoannulata***

7b. Basidiomata with salmon pinkish tinges and paler, more fragile and more spaced lamellae; terminal cells of the pileipellis predominantly fusiform to lageniform. Locally common.

***Russula cameroonensis***

## Acknowledgements

The first author thanks Terence Fuh and the staff of the Primate Habituation Programme of the Dzanga-Ndoki National Park, “Réserve Spéciale de Foret Dense de Dzanga-Sangha” at Bayanga, as well as all staff, Aka guides and visitors of the Bai Hokou field station for their logistical support, field assistance and their very enjoyable company. This research was made possible through research permit 034/MENESR/DIRCAB/DGESRSTI/DRSTSPI/SSSTI/16 from the “Ministère de l’éducation nationale, de l’enseignement supérieur et de la recherche scientifique” of the Central African Republic. In Cameroon, the Ministry of Research and Scientific Innovation issued research permits. The Conservator of the Dja Biosphere Reserve, Mr. Ndinga Hilaire, and his staff greatly assisted the fieldwork in the Dja. Staff of the Congo Basin Institute provided much logistical support. Field assistance in Cameroon was provided by Mei Lin Chin, Todd Elliott, Camille Truong, Bryn Dentinger, Cathie Aime, Rachel Koch, Blaise Jumbam, Olivier Séné, Carolyn Delevich, Kennan Mighell, Jessie Uehling, Noah Siegel, Alamane Gabriel (a.k.a. Sikiro), and Essambe Jean-Pierre (a.k.a. Papa Chef). The first author thanks the Curator of the Herbarium (BR) of the Botanic Garden Meise, Belgium, for permission to reproduce the original watercolors by Mme. Goossens-Fontana, and M. Härkönen and T. Niemelä for providing info and photograph for *R. acriannulata*. Cathrin Manz thanks Felix Hampe, Daouda Dongnima and Gildas Abouhoumbo for assistance in the field, further also Meike Piepenbring and Nourou S. Yorou for coordination of the project FunTrAf. and Niklas Döring is thanked for technical support with scanning electron microscopy. Dr. Karen Hansen, Swedish Museum of Natural History, W.B Li of Kunming Institute of Botany, Chinese Academy of Sciences, helped with the field trips in Sweden and China and lab work. Herbarium GDGM, Guandong, China genereously provided loan of holotype of *R. verrucospora* for our study.

